# piRNAs are regulators of metabolic reprogramming in stem cells

**DOI:** 10.1101/2023.10.31.564965

**Authors:** Patricia Rojas-Ríos, Aymeric Chartier, Camille Enjolras, Julie Cremaschi, Céline Garret, Adel Boughlita, Anne Ramat, Martine Simonelig

## Abstract

Stem cells preferentially use glycolysis instead of oxidative phosphorylation and this metabolic rewiring plays an instructive role in their fate; however, the underlying molecular mechanisms remain largely unexplored. PIWI-interacting RNAs (piRNAs) and PIWI proteins have essential functions in a range of adult stem cells across species. Here, we show that piRNAs and the PIWI protein Aubergine (Aub) are instrumental in activating glycolysis in *Drosophila* germline stem cells (GSCs). High glycolysis is required for GSC self-renewal and *aub* loss-of-function induces a metabolic switch in GSCs leading to their differentiation. Aub directly binds glycolytic mRNAs and *Enolase* mRNA regulation by Aub depends on its 5’UTR. Furthermore, deletion of a piRNA target site in *Enolase* 5’UTR leads to GSC loss. These data reveal an Aub/piRNA function in translational activation of glycolytic mRNAs in GSCs, and pinpoint a new mode of regulation of metabolic reprogramming in stem cells based on small RNAs.

## Introduction

Energy metabolism has long been known as a homeostatic system that can adapt to cellular energy needs. However, a growing body of evidences have now directly implicated energy metabolism in cell fate. Metabolic remodeling takes place early during transitions between cell states, before the establishment of a particular cell fate, and has been shown to play an important role in stem cell biology and reprogramming. In particular, induction of pluripotency during cell reprogramming correlates with a shift from oxidative phosphorylation (oxphos) to glycolysis, and stimulation of glycolysis facilitates pluripotency ^1, 2^. Conversely, oxidative metabolism is activated during the differentiation of pluripotent stem cells, and differentiation can be suppressed through decreased mitochondrial respiration ^1, 3, 4^. These findings underscore the importance of energetic pathways in pluripotency and stem cell differentiation. Increased glycolysis is a common feature across a number of stem cell populations, including hematopoietic and mesenchymal stem cells ^5–7^. One reason of the glycolytic state of stem cells is to preserve their genome integrity by maintaining low oxidative stress through reduced oxphos ^8^. A key question is thus to understand how energy metabolism is controlled in stem cells at the molecular level. Regulation occurs in part at the transcriptional level through hypoxia as various stem cell populations reside within niches with low oxygen tension. The transcription factors hypoxia-inducible factor (HIF)1α and 2β are stabilized under hypoxic conditions and associate with a constitutive HIF subunit to promote glycolysis and suppress mitochondrial biogenesis. This occurs through transcriptional activation of several glycolytic genes and repression of PGC-1α, a general regulator of genes encoding mitochondrial proteins ^9, 10^. Besides this transcriptional regulation, very little is known regarding the regulatory mechanisms underlying metabolic remodeling in stem cells, despite the importance of this remodeling in stem cell fate.

Piwi-interacting RNAs (piRNAs) are a class of small non-coding RNAs first identified in the germline of animal species, but widely present in somatic tissues as well ^11^. piRNAs are loaded into specific Argonaute proteins, called PIWI proteins, and interact with their target mRNAs by complementarity. piRNAs and PIWI proteins are involved in repressing transposable elements, but they also have essential functions as regulators of cellular mRNAs *via* various mechanisms. These include mRNA cleavage through the endonuclease activity of PIWI proteins, deadenylation and decay, stabilization, and translational activation, thanks to the capacity of PIWI proteins to recruit different mRNA regulatory machineries ^12, 13^. PIWI proteins and piRNAs are required in various stem cell populations. *Drosophila* Piwi, the founding member of the PIWI family was identified based on its function in germline stem cell (GSC) asymmetric division ^14^. Piwi is necessary in both GSCs and somatic niche cells for GSC self-renewal and differentiation ^15^. The functions of PIWI proteins in GSC proliferation and maintenance are conserved from *C. elegans* to the mouse ^16, 17^. Recently, a role of Aubergine (Aub), another of the three *Drosophila* PIWI proteins was also demonstrated in GSC self-renewal and to a lesser extent differentiation ^18, 19^. In *aub* mutant, GSCs can differentiate into germ cells, but they lose the capacity to self-renew and are rapidly lost. This role of Aub depends on its capacity to bind piRNAs and its function in gene regulation ^19^. The proto-oncogene *Cbl* was identified as a relevant Aub mRNA target, and Aub acts as a negative regulator of *Cbl* mRNA for GSC self-renewal ^19^. However, Aub positively regulates other mRNAs such as *dunce*, and was proposed to do so through activation of translation initiation ^18^.

PIWI proteins are also required in stem cell populations outside the gonads. *Drosophila* Piwi has functions in the gut in establishing the intestinal stem cell population in pre-adult stages ^20^, and in the maintenance and differentiation of intestinal stem cells in response to stress and during aging ^21^. Strikingly, PIWI proteins are also markers of pluripotent stem cells in primitive species with high regenerative capacities, such as sponges, cnidarians and planarians ^22–24^. Moreover, knock-down approaches have revealed their requirement for whole-body regeneration in several of these species^15, 23^.

Our understanding of piRNA biogenesis has revealed tight links between energy metabolism and the piRNA pathway. piRNA biogenesis occurs in part at the mitochondrial outer membrane and the endonuclease (*Drosophila* Zucchini, mouse MitoPLD), as well as other factors involved in piRNA production are integral components of the mitochondrial outer membrane ^25–28^. In addition, a genetic screen identified a role of subunits of the mitochondrial respiratory chain complexes in piRNA biogenesis ^29^. Moreover, glycolytic enzymes were also reported to be involved in piRNA biogenesis ^30^. Using Aub iCLIP (individual-nucleotide resolution UV crosslinking and immunoprecipitation) datasets from early embryos ^31^ and cultured GSCs ^18^, we identified a total of seven glycolytic mRNAs among mRNAs encoding the eleven glycolytic enzymes (Figure 1A). This suggested that Aub could regulate glycolysis through direct interaction with glycolytic mRNAs. Here, we show that glycolysis is the main energetic pathway in GSCs and that high glycolysis is required for GSC self-renewal. Either reducing glycolysis by decreasing the levels of glycolytic enzymes, or increasing oxphos in GSCs promote GSC differentiation, preventing their maintenance. Strikingly, a switch in energy metabolism towards oxphos takes place in *aub* mutant GSCs. Aub-dependent increase of glycolytic enzyme levels in GSCs compared to differentiating cells, together with piRNA-dependent Aub binding to glycolytic mRNAs suggest a direct role of Aub in translational activation of glycolytic mRNAs in GSCs. This was confirmed using reporter transgenes containing the 5’UTR of *Enolase* (*Eno*) that were found to display Aub-dependent higher expression in GSCs. In addition, CRISPR-based deletions of a piRNA target site in *Eno* 5’UTR results in GSC loss. Finally, the strong GSC loss characteristic of *aub* mutants was reduced by increasing glycolysis. Together, these results reveal a key role of Aub and piRNAs in activating glycolysis in GSCs for their self-renewal. They also identify a new regulatory mechanism of the metabolic remodeling in stem cells depending on piRNAs. Because piRNAs and PIWI proteins are present in various stem cell populations including in primitive regenerative species, this mode of regulation of stem cell metabolic reprogramming might be conserved throughout evolution.

**Figure 1.**
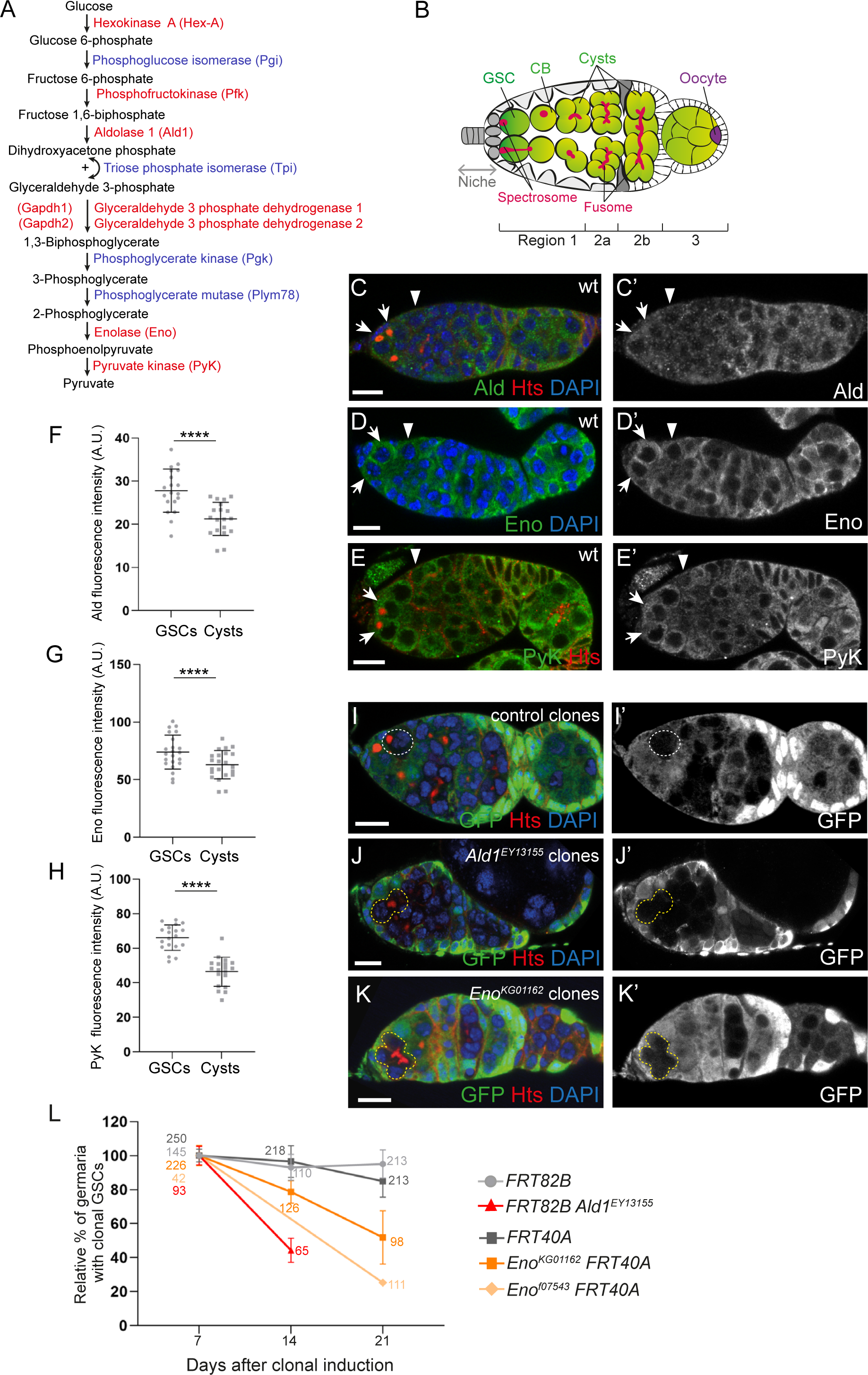
Glycolytic genes are required for GSC self-renewal. (A) The ten steps of glycolysis. Enzymes whose mRNAs were found in GFP-Aub iCLIP datasets ^18, 31^ are in red. Aub binding to glycolytic mRNAs is statistically significant, *p*=1.0e-05 using the Generalized hypergeometric test. (B) Schematic diagram of the germarium of the *Drosophila* ovary. Germ cells are in green, except the oocyte that is in purple (GSC: germline stem cell; CB: cystoblast). The spectrosome (spherical) and fusome (branched) are in red. During GSC division, the spectrosome acquires an elongated form in the shape of an exclamation mark. Differentiation proceeds from anterior (left) to posterior (right). Three regions have been described in the germarium: regions 1, 2 and 3 containing mitotically active germ cells, differentiating 16 cell-cysts, and the newly formed egg chamber, respectively. (C-E’) Confocal images of wild-type (wt) germaria immunostained with an anti-Ald (C, C’), anti-Eno (D, D’) or anti-PyK (E, E’) (green), anti-Hts (red) and DAPI. White arrows indicate GCSs and white arrowheads indicate the region containing germline cysts where glycolytic enzyme levels were quantified. (F-H) Quantification of Ald1 (F), Eno (G) and PyK (H) protein levels in GSCs and differentiating cyst cells using immunostaining experiments shown in C-E’. Fluorescence intensity was measured in arbitrary units using the ImageJ software. Horizontal bars represent the mean and error bars represent standard deviations. \*\*\*\**p*-value <0.0001 using the paired Student’s *t*-test. (I-K’) Confocal images of mosaic germaria containing control (I, I’), *Ald1^EY13155^* (J, J’) or *Eno^KG01162^* (K, K’) clonal cells stained with anti-GFP (green) (lack of GFP indicates clonal cells), anti-Hts (red) and DAPI (blue), 14 days after clonal induction. The white dashed line (I, I’) indicates a control clonal GSC, and the yellow dashed line (J-K’) indicate mutant clonal differentiating cysts in contact with the niche. Clone induction was performed in adults to analyze to role of *Ald1* and *Eno* in GSCs during adult stages. Heterozygous *Ald1^EY13155^* females in which mitotic clones have been induced are not viable after 17 days. (L) Relative percentages of germaria with at least a clonal GSC at 7, 14 or 21 days after clonal induction. The number of germaria analyzed is indicated. Error bars represent standard deviations. Scale bars: 10 μm.

## Results

### High glycolysis is required for GSC self-renewal

Although various populations of stem cells are known to preferentially use glycolysis as their main energetic pathway, this question has not been addressed in *Drosophila* ovarian GSCs, one of the best-described model of adult stem cells. The *Drosophila* ovary is composed of 16 to 18 ovarioles in which egg chambers develop towards the posterior. The anterior-most region of ovarioles is called the germarium and contains two to three GSCs localized anteriorly and in contact with the niche composed of somatic cells. GSCs are identified by a spherical organelle, the spectrosome localized anteriorly towards the niche (Figure 1B). GSCs divide asymmetrically to produce a new GSC -the cell that remains in contact with the niche-by a process known as self-renewal, and a cystoblast -the cell that loses the contact with the niche-that starts differentiation. The cystoblast divides four consecutive times with incomplete cytokinesis to produce a germline cyst composed of sixteen interconnected germline cells containing a branched fusome derived from the original spectrosome (Figure 1B).

To document the metabolic reprogramming in *Drosophila* GSCs, we first analyzed the expression patterns of three glycolytic enzymes, Aldolase 1 (Ald1), Eno and Pyruvate kinase (PyK) in germaria using immunostaining. We used available antibodies directed against mammalian proteins since these proteins are highly conserved. The three glycolytic enzymes were present in all cells in the germarium consistent with their fundamental role in energy metabolism, but their levels were higher in GSCs than in early differentiating germ cells (Figures 1C-1H). The specificity of the three antibodies was confirmed by immunostaining of germaria expressing the corresponding RNAi with the *UAS/Gal4* system and the germline-specific driver *nos-Gal4* (Figures S1A-S1F). PyK higher protein levels in GSCs compared to differentiating germ cells was validated using a *P-GFP* (*Wee-P*) insertion in the *PyK* locus producing a GFP-PyK fusion protein ^32^ (Figures S1G-S1H). Consistent with these data, previous studies have shown that GSCs do not express high levels of oxphos components and have reduced mitochondrial membrane potential ^33, 34^. Specifically, subunits of ATP synthase, the last complex of the mitochondrial electron transport chain were not found in GSCs, but were present in high amounts in differentiating cyst cells ^33^. In addition, mitochondrial membrane potential as well as electron transport chain activity were very low in GSCs and the following dividing cyst cells and sharply increased in sixteen cell-cysts ^34^.

We next determined whether glycolysis plays an instructive role in GSC stemness. We addressed the requirement of high glycolysis for GSC self-renewal through reduction of glycolytic enzyme levels using clonal analysis with *Eno* and *Ald1* mutants. The *Eno^KG601162^* and *Ald1^EY13155^* ^35^ (Figure S1I) mutants are lethal but clonal mutant germ cells do survive, suggesting their hypomorphic nature ^30^. Wild-type, *Ald1^EY13155^* and *Eno^KG601162^* clonal GSCs were generated with the FLP-mediated FRT recombination system ^36^ and the presence of clonal GSCs, revealed by the loss of GFP expression, was quantified at three time points, i.e. 7, 14 and 21 days after clone induction. Wild-type clonal GSCs were maintained over time (Figures 1I, 1I’, 1L), whereas a proportion of *Ald1^EY13155^* and *Eno^KG601162^* mutant GSCs were lost 14 and 21 days after clone induction, showing a defect in their self-renewal (Figures 1J-1K’, 1L). Importantly, immunostaining of these germaria with anti-Hts antibody that marks spectrosomes and fusomes showed that both *Ald1* and *Eno* mutant differentiating cysts were present in the niche, indicating that mutant clonal GSCs were able to differentiate and were likely lost by differentiation (Figures 1J, 1K). This was confirmed by recording cell death in clonal mutant GSCs using anti-cleaved Caspase 3 staining. The percentage of germaria with clonal GSCs positive for anti-cleaved Caspase 3 was very low and not higher in mutant than wild-type GSCs (Figures S1J-S1N), showing that *Ald1* and *Eno* mutant GSCs did not undergo apoptosis, but were lost by differentiation.

To confirm the role of glycolytic genes in GSC self-renewal, we expressed available *UAS-RNAi* transgenes with the *nos-Gal4* driver to reduce the levels of glycolytic enzymes in GSCs. *UAS-RNAi* transgene efficiency was validated using RT-qPCR (Figure S2A). Immunostaining of ovaries with anti-Hts antibody, 7, 14 and 21 days after eclosion, showed that downregulation in germ cells of the nine tested glycolytic mRNAs led to GSC loss quantified by the percentage of germaria with less than two GSCs (Figures S2B-S2L). This loss increased with time and differentiated cysts could be found in the niche, indicating that differentiation was not prevented.

These results show that glycolytic enzymes accumulate at higher levels in GSCs than in differentiating cysts and that high levels of glycolytic enzymes are required for GSC self-renewal.

### Low Oxphos is required for GSC self-renewal

In addition to the increased levels of ATP synthase components in differentiating cysts, these components were shown to be required for division and differentiation of cyst cells, but not for GSC self-renewal ^33^. ATP synthase is involved in the maturation of mitochondrial cristae in differentiating cyst cells, a prerequisite for oxphos ^33^. To strengthen the conclusion of metabolic reprogramming towards glycolysis playing an important role in GSC homeostasis, we analyzed the effect of increasing oxphos in GSCs using two different approaches. First, we deregulated the mitochondrial pyruvate dehydrogenase (PDH) complex that catalyzes the first step of pyruvate metabolism for oxphos (Figure 2A). The E1 enzyme of PDH is activated by pyruvate dehydrogenase phosphatase (PDP) and inhibited by pyruvate dehydrogenase kinase (PDK) ^37, 38^. We used the *nos-Gal4* driver to express *UAS-RNAi* transgenes in germ cells and reduce the levels of *PDP* or *PDK* mRNAs (Figures S2M, S2N). Reducing PDP levels should reduce oxphos and this had no effect on GSC self-renewal, consistent with GSCs not heavily depending on oxphos (Figures 2B, 2C, 2G). In contrast, reducing PDK levels that was expected to favor oxphos, led to GSC loss visualized by the presence of differentiating cysts in the niche, which increased with time (Figures 2D, 2G). Next, we increased oxphos in germ cells by overexpressing general activators of mitochondrial mass and activity. The transcription factor nuclear respiratory factor-1 (NRF-1, Erect wing in *Drosophila*) and the coactivator PPAR coactivator 1 (PGC-1, Spargel in *Drosophila*) coregulate a large set of genes involved in mitochondrial function ^39–41^. *erect wing* (*ewg*) and *spargel* were overexpressed in germ cells using *P-UAS* constructs inserted in the genes and the *nos-Gal4* driver. Overexpression of both *ewg* and *spargel* (Figures S2O, S2P) led to GSC loss and the presence of differentiating cyst cells in the niche, showing that increased oxphos prevented GSC self-renewal, but not differentiation (Figures 2E-2G).

**Figure 2.**
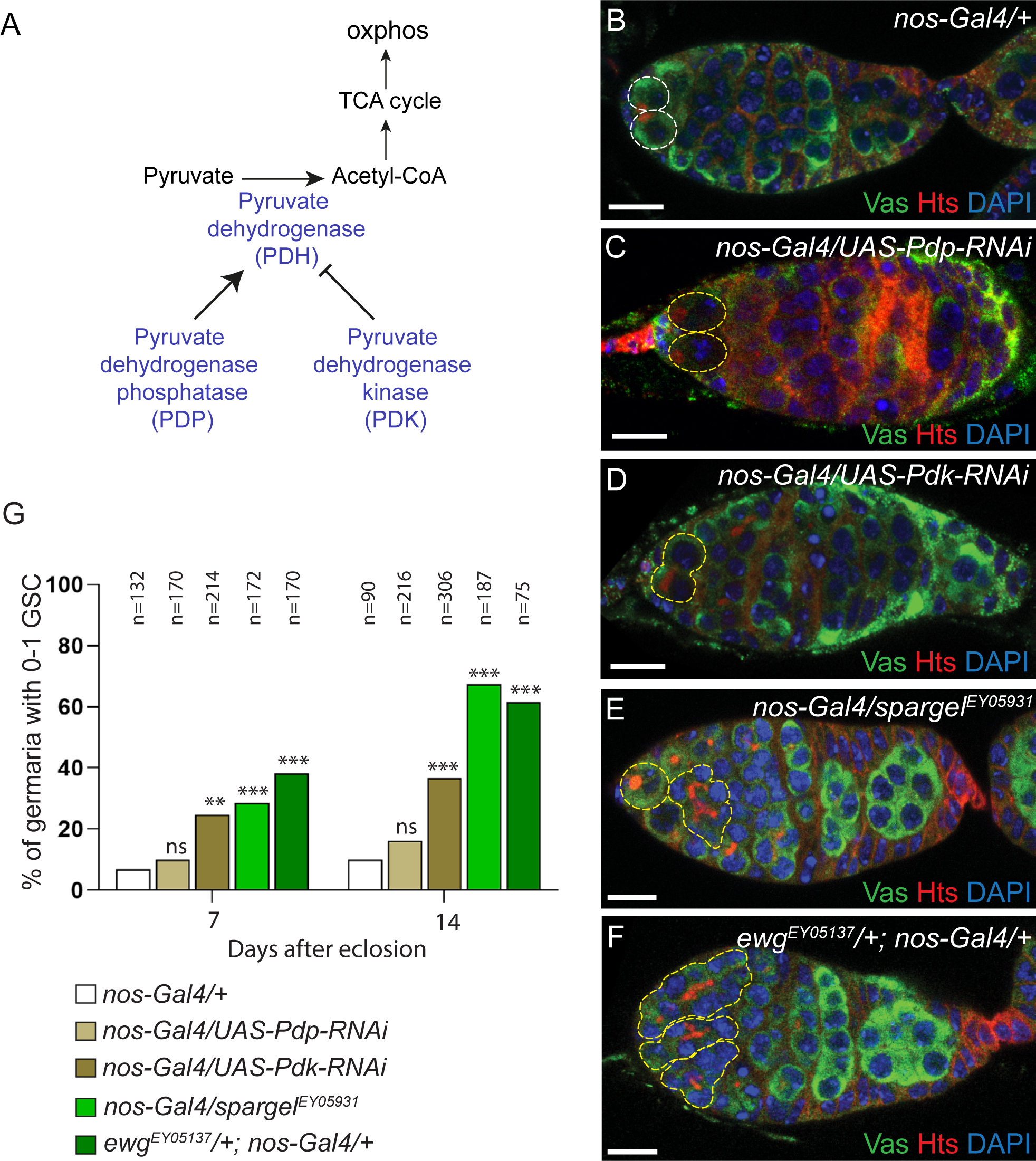
Increased mitochondrial activity in germ cells leads to GSC loss. (A) Scheme of Pyruvate Dehydrogenase (PDH) complex regulation by Pyruvate Dehydrogenase Phosphatase (PDP) and Pyruvate Dehydrogenase Kinase (PDK). (B-F) Confocal images of germaria from 7 day old-females stained with anti-Vasa (green), anti-Hts (red) and DAPI (blue). The white dashed line indicates two GSCs in a control *nos-Gal4/+* germarium (B) and the yellow dashed lines indicate two GSCs in a *nos-Gal4/UAS-Pdp-RNAi* germarium (C) and a single GSC or differentiating cysts in the niche in *nos-Gal4/UAS-Pdk-RNAi* (D), *nos-Gal4/spargel^EY05931^* (E) and *ewg^EY05137^/+; nos-Gal4/+* (F) germaria, respectively. (G) Quantification of germaria showing GSC loss (0-1 GSC) 7 and 14 days after eclosion. The number of scored germaria (n) is indicated. \*\**p*-value <0.01, \*\*\**p*-value <0.001, ns, non-significant using the χ^2^ test. Scale bars: 10 μm.

These data reveal that high oxphos levels are not compatible with GSC maintenance. They are consistent with a recent study showing that increasing mitochondrial membrane potential by interfering with mitochondrial dynamics results in GSC loss ^42^.

### Aubergine is required for metabolic reprogramming towards glycolysis in GSCs

Aub iCLIP assays revealed that glycolytic mRNAs interact with Aub. We, therefore, asked whether Aub could modulate energy metabolism by directly regulating glycolytic mRNAs. We first performed immunostaining of *aub* mutant ovaries with anti-Eno and anti-PyK antibodies to address whether the higher levels of glycolytic enzymes found in wild-type GSCs were maintained in *aub* mutant GSCs. Both Eno and PyK levels were similar in GSCs and differentiating cyst cells, in *aub* mutant germaria (Figures 3A-3E’), as quantified by the ratio of immunofluorescence intensity in GSCs to cyst cells that was significantly reduced in *aub* mutant compared to wild-type germaria (Figures 3C, 3F). Thus, Aub increases glycolytic enzyme levels in GSCs, possibly through direct regulation of glycolytic mRNAs.

**Figure 3.**
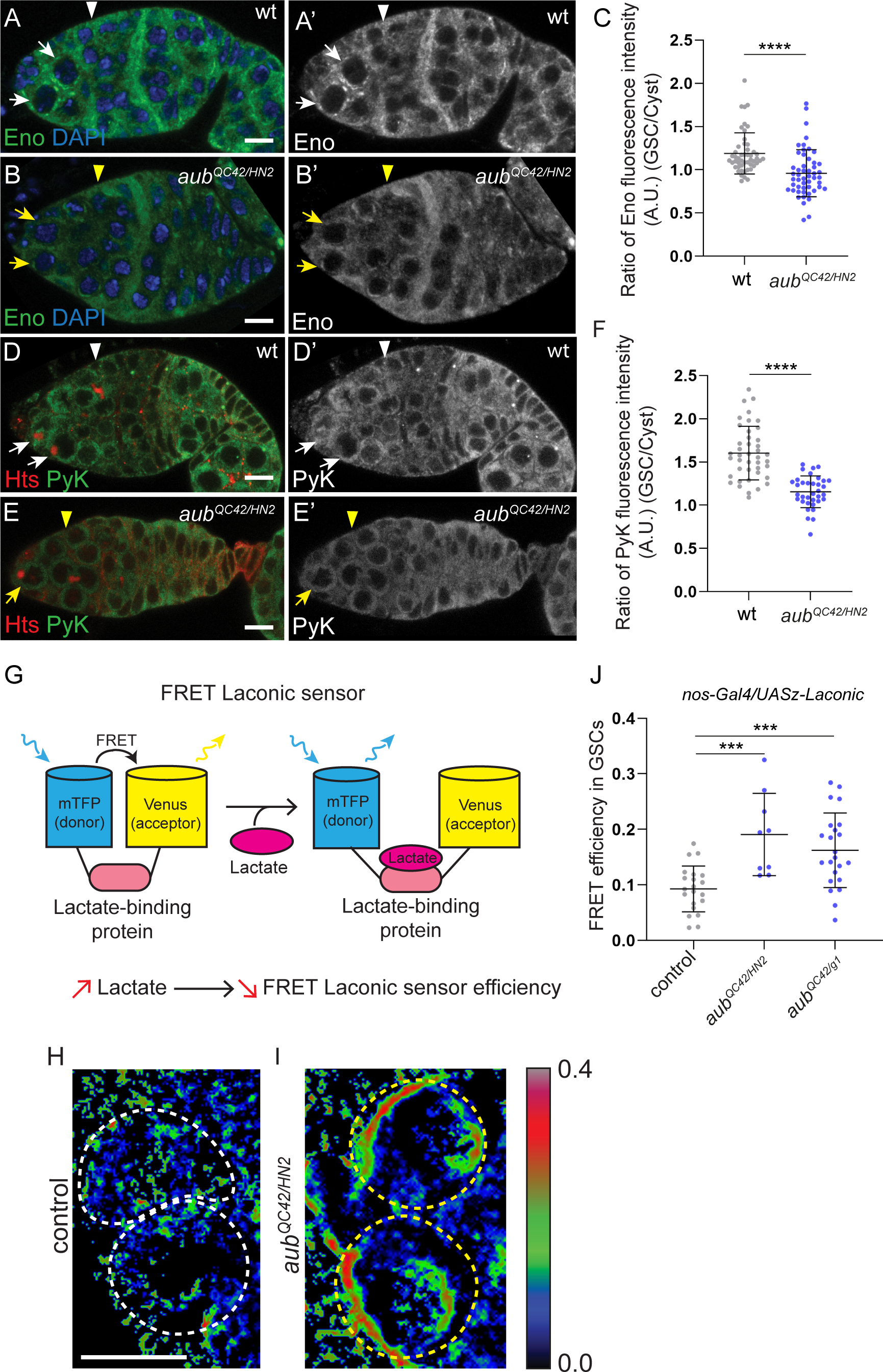
Aub increases glycolytic enzymes and lactate levels in GSCs. (A-B’, D-E’) Confocal images of immunostaining of wt and *aub^QC42/HN2^* germaria with anti-Eno (green) and DAPI (blue) (A-B’) or anti-PyK (green) and anti-Hts (red) (D-E’). White and yellow arrows indicate GCSs in wt and *aub* mutant germaria, respectively. White and yellow arrowheads indicate the region containing germline cysts where glycolytic enzyme levels were quantified, in wt and *aub* mutant germaria, respectively. (C, F) Quantification of Eno and PyK protein levels in wt and *aub^QC42/HN2^*mutant GSCs and differentiating cyst cells using immunostaining experiments shown in (A-B’, D-E’). Fluorescence intensity was measured in arbitrary units using the ImageJ software. Ratios of fluorescence intensity of one GSC to one cyst cell per germarium were plotted. Horizontal bars represent the mean and error bars represent standard deviations. \*\*\*\**p*-value <0.0001 using the unpaired Student’s *t*-test. (G) Schematic representation of the FRET Laconic sensor. The binding of lactate to the ligand-binding domain leads to a conformational change that increases the distance between the donor (mTFP) and the acceptor (Venus), resulting in reduced FRET efficiency. Thus, FRET Laconic sensor efficiency inversely correlates with lactate concentration. (H, I) FRET ratio images in control *nos-Gal4/UASz-Laconic* (H) and *aub* mutant, *aub^QC42/HN2^; nos-Gal4/UASz-Laconic* (I) anterior-most region of germaria. Control and *aub* mutant GSCs are indicated with white and yellow dashed lines, respectively. The rainbow colormap indicates the FRET efficiency levels. (J) Quantification of FRET Laconic sensor efficiency in control and *aub* mutant, *aub^QC42/HN2^* and *aub^QC42/g1^*, GSCs from three day old-females based on acceptor photobleaching. \*\*\**p*-value <0.001 using the unpaired Student’s *t*-test. Scale bars: 10 μm in A-E’; 5 μm in H, I.

To more directly assess the role of Aub in the regulation of energy metabolism in GSCs, we implemented the utilization of FRET metabolic sensors in GSCs. These genetically-encoded fluorescent probes sensitive to intracellular concentration of metabolites allow to quantify specific metabolites at single cell resolution. Pyruvate, the end-product of glycolysis is the input energy substrate for oxphos and can be converted into lactate under low oxphos. We used a lactate sensor called Laconic (an *UASz* version of the Laconic sensor ^43, 44^) expressed in germ cells with the *nos-Gal4* driver. This Laconic sensor is built such that FRET efficiency decreases in the presence of lactate (Figure 3G). FRET efficiency of the Laconic sensor was quantified based on acceptor photobleaching (see Methods). We first validated that the Venus acceptor alone did not produce donor mTFP-like fluorescence by photoconversion upon photobleaching ^45^ (Figures S3A-S3B’). FRET Laconic sensor efficiency increased in *aub* mutant GSCs compared to wild-type revealing lower lactate levels (Figures 3H-3J). These reduced amounts of lactate indicated lower glycolysis and/or higher oxphos in *aub* mutant GSCs, consistent with the proposed role of Aub in activating glycolysis in GSCs.

We next addressed whether mitochondrial morphology and activity might be affected in *aub* mutant GSCs. Mitochondrial activity is intimately linked to their structure and organization. Globular fissed mitochondria with underdeveloped cristae are a hallmark of stem cells and induced pluripotent stem cells that mainly rely on glycolysis, whereas fused mitochondria with numerous cristae are present in differentiated cells allowing ATP production through increased oxphos activity ^8, 46–48^. Importantly, in *Drosophila* GSCs, mitochondria were also reported to be fissed with poorly developed cristae, and they became more fused with abundant cristae following differentiation ^33, 42^. We thus analyzed mitochondrial morphology in wild-type and *aub* mutant GSCs using transmission electron microscopy. As described previously, in wild-type GSCs mitochondria tended to be globular with undeveloped cristae (Figures 4A, 4A’). In *aub* mutant GSCs, mitochondria were more fused as shown by the increased number of longer mitochondria compared to their length in wild-type GSCs, and they had more developed cristae (Figures 4B-4D). Therefore, in *aub* mutant GSCs, the mitochondrial network was more mature than in wild-type GSCs, resembling that in differentiating cells. We then analyzed the expression of ATP synthase α (ATPsynα) in *aub* mutant GSCs. In wild-type GSCs, the levels of ATPsynα are very low in GSCs and strongly increase in differentiating 8- and 16-cell cysts ^33^. We recorded ATPsynα levels in *aub* mutant GSCs by inducing *aub* mutant clonal GSCs followed by immunostaining of ovaries with anti-ATPsynα antibody. *aub* mutant GSCs expressed significantly higher levels of ATPsynα than control heterozygous GSCs (Figures 4E-4F). To investigate whether reduced glycolysis in GSCs could lead to indirect increased expression of oxphos genes, we quantified ATPsynα expression in *Ald1* and *Eno* mutant GSCs using the clonal analysis. Similarly to *aub* mutant GSCs, the levels of ATPsynα were higher in *Ald1* and *Eno* mutant GSCs than in control heterozygous GSCs, suggesting a metabolic rewiring upon lower glycolysis (Figures S4A-S4F).

**Figure 4.**
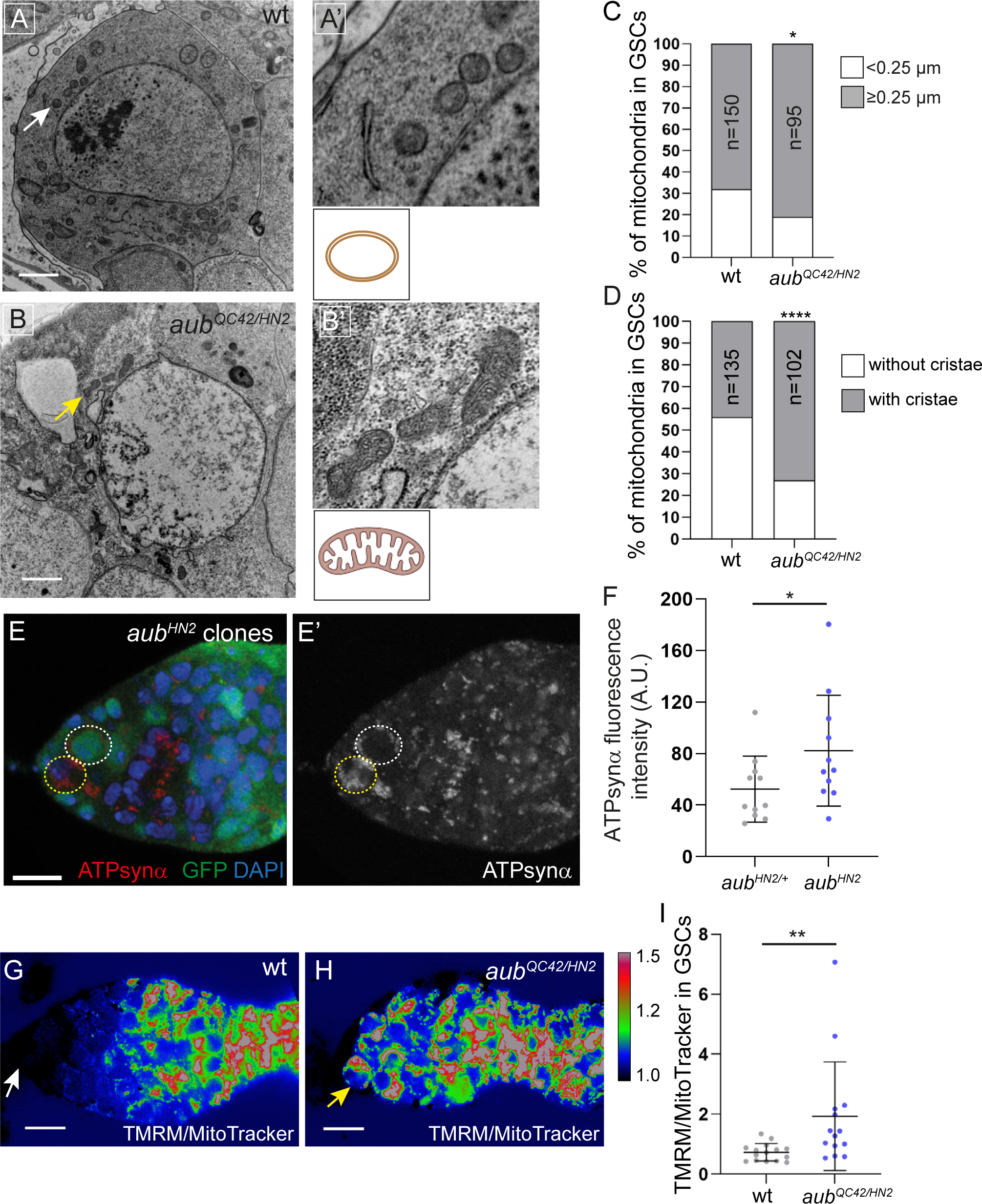
The lack of Aub induces a switch in energy metabolism in GSCs. (A-B’) Transmission Electron Microscopy images of a wt (A) and an *aub^QC42/HN2^*mutant (B) GSC. The white arrow in wt and yellow arrow in *aub* mutant GSCs point to mitochondria that are shown enlarged in (A’) and (B’). (C) Measurement of mitochondrial length in wt and *aub^QC42/HN2^* mutant GSCs from the Transmission Electron micrographs. Percentages of mitochondria with a length above or below 0.25 μm are shown. The number of mitochondria (n) from at least ten GSCs per condition is indicated. \**p*-value <0.05 using the χ^2^ test. (D) Quantification of mitochondria with and without cristae in wt and *aub^QC42/HN2^* mutant GSCs from the Transmission Electron micrographs. Percentages of mitochondria with and without cristae are shown. The number of mitochondria (n) from at least ten GSCs per condition is indicated. \*\*\*\**p*-value <0.0001 using the χ^2^ test. (E, E’) Confocal images of a mosaic germarium containing an *aub^HN2^* mutant clonal GSC stained with anti-GFP (green), anti-ATPsynα (red) and DAPI (blue). The non-clonal (GFP^+^) and *aub* mutant clonal (GFP^-^) GSCs are indicated by dashed white and yellow lines, respectively. (F) Quantification of ATPsynα protein levels in control non-clonal *aub^HN2/+^* and clonal *aub^HN2^* mutant GSCs using fluorescence intensity of immunostaining with anti-ATPsynα. Fluorescence intensity was measured in arbitrary units using the ImageJ software. Horizontal bars represent the mean and error bars represent standard deviations. \**p*-value <0.05 using the paired Student’s *t*-test. (G, H) Quantification of mitochondrial membrane potential: ratiometric images of wt and *aub^QC42/HN2^* mutant germaria stained with TMRM to record mitochondrial membrane potential and Deep Red MitoTracker to record mitochondrial mass. The ratio of TMRM to MitoTracker intensity was calculated using ImageJ and a pseudo-colored image was generated using the rainbow RGB gradient. White and yellow arrows indicate wt and *aub* mutant GSCs, respectively. (I) Quantification of TMRM/MitoTracker ratios in wt and *aub^QC42/HN2^* mutant GSCs using ImageJ. ** *p*-value <0.01 using the Mann-Whitney test. Scale bars: 1 μm in (A, B); 10 μm in E, G, H.

Finally, we quantified mitochondrial membrane potential as a measurement of mitochondrial activity in wild-type and *aub* mutant GSCs, using tetramethylrhodamine methyl ester (TMRM) staining to record mitochondrial membrane potential and Deep Red MitoTracker to measure mitochondrial mass ^34, 42^. As reported previously, in wild-type germaria mitochondrial membrane potential was very low in GSCs and dividing cyst cells, and sharply increased in 16 cell-cysts (Figures 4G, 4I) ^34^. In contrast, in *aub* mutant germaria mitochondrial membrane potential was already high in GSCs (Figures 4H, 4I).

Together these results show that Aub plays a key role in increasing glycolysis in GSCs. In the absence of Aub, expression of glycolytic enzymes is reduced in GSCs and this is accompanied by a switch in energy metabolism towards oxphos with higher expression of ATP synthase that might contribute to mitochondrial maturation.

### Aubergine directly regulates glycolytic mRNAs

GFP-Aub iCLIP assays performed in cultured GSCs revealed five glycolytic mRNAs significantly bound by Aub ^18^ (Figure S5A). Aub iCLIP sites were mostly found in 5’UTRs and 3’UTRs, with higher accumulation in 3’UTRs in most cases. To confirm Aub interaction with glycolytic mRNAs and address whether this interaction depends on Aub capacity to load piRNAs, we performed RNA immunoprecipitation (RIP) in ovaries. GFP-Aub and GFP-Aub^AA^, an Aub double point mutant in the PAZ domain preventing Aub loading with piRNAs were expressed in GSCs and germ cells using *UASp*-based transgenes and the *nos-Gal4* driver ^31^ (Figures 5A, 5B). GFP-Aub and GFP-Aub^AA^ immunoprecipitations were validated using western blots (Figure 5C) and enrichment of glycolytic mRNAs in GFP-Aub and GFP-Aub^AA^ RIPs were quantified using RT-qPCR (Figures 5D, 5E, S5B). The seven glycolytic mRNAs identified in GFP-Aub iCLIP data sets from GSCs and embryos ^18, 31^ were enriched in GFP-Aub RIPs, but not in GFP-Aub^AA^ RIPs (Figures 5D, 5E, S5B), confirming Aub interaction with these mRNAs and demonstrating the requirement of Aub loading with piRNAs for this interaction.

**Figure 5.**
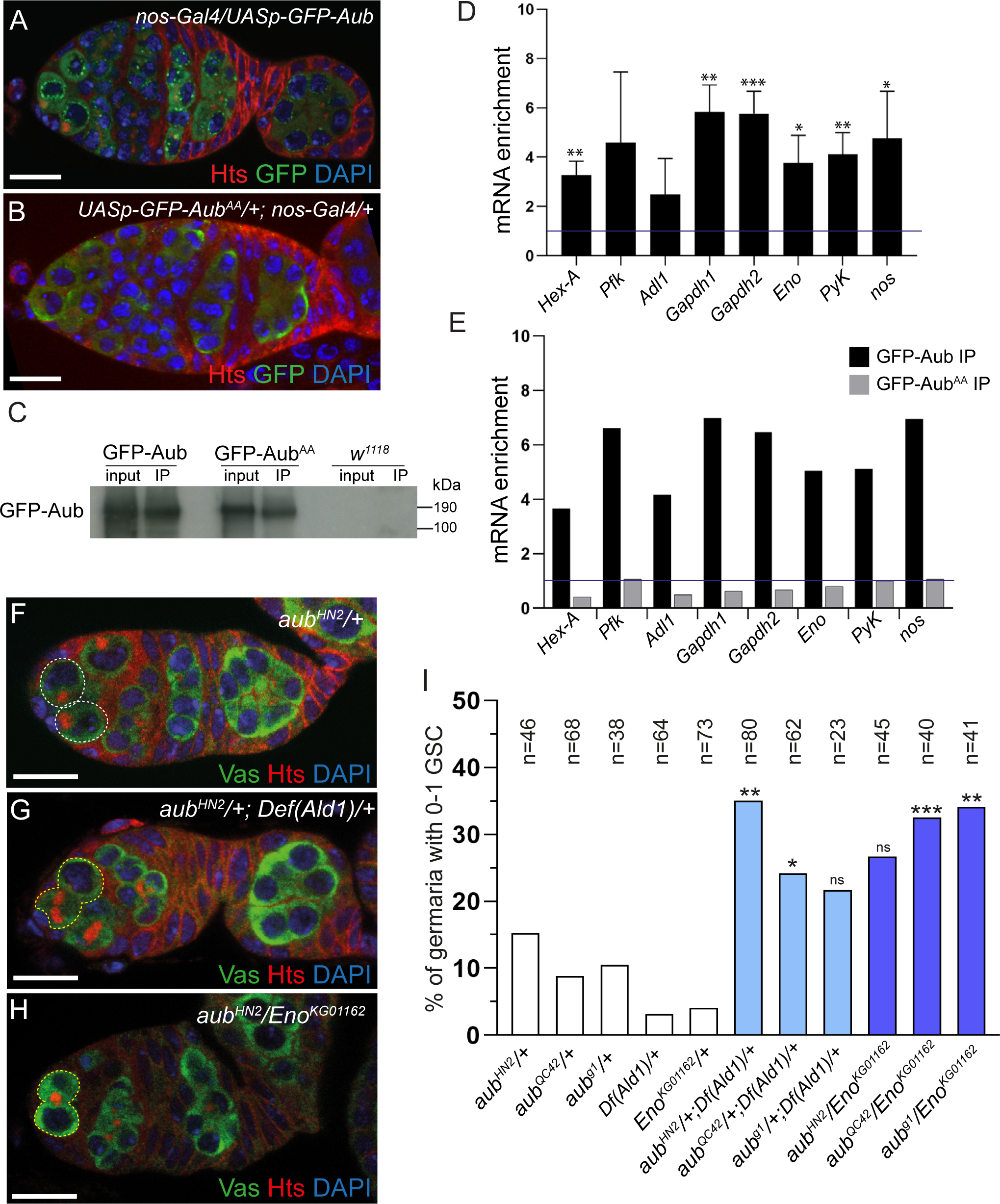
Aub binds to and activates glycolytic mRNAs. (A, B) Confocal images of immunostaining of germaria expressing GFP-Aub (*nos-Gal4/UASp-GFP-Aub*) or GFP fused to the double point Aub mutant that does not load piRNAs, Aub^AA^ (*UASp-GFP-Aub*^AA^/+; *nos-Gal4/+*) with anti-GFP (green), anti-Hts (red) and DAPI (blue). GFP-Aub is present in the nuage around nuclei, whereas GFP-Aub^AA^ is more diffuse and not present in the nuage. (C) Western blot showing immunoprecipitation (IP) of GFP-Aub and GFP-Aub^AA^ with anti-GFP in ovaries. Genotypes were as in (A, B). *w^1118^* ovaries were used as a negative control. Input corresponds to 1/8 of the extract prior to immunoprecipitation. The blot was revealed with anti-GFP. (D, E) Quantification of mRNAs using RT-qPCR in GFP-Aub (D, E) and GFP-Aub^AA^ (E) IPs. mRNA enrichment compared to mRNA levels in *w^1118^* IPs set to 1 (horizontal bars). mRNA levels were normalized to U1 snRNA and to RNA levels in the input. *nanos* (*nos*) mRNA was used as a positive control ^31^. Mean of three biological replicates in (D). Error bars represent standard deviation. * *p*- value <0.05, \*\**p*-value <0.01, *** *p*-value <0.001 using the unpaired Student’s *t*-test. One biological replicate in (E) showing the lack of mRNA enrichment in GFP-Aub^AA^ IP. (F-I) Genetic interaction between *aub* and either *Ald1* or *Eno* for GSC self-renewal. Confocal images of immunostaining of *aub^HN2^/+* (F), *aub^HN2^/+*; *Def(Ald1)*/+ (G) and *aub^HN2^/Eno^KG01162^* (H) germaria with anti-Vasa (green), anti-Hts (red) and DAPI (blue). The white dashed line indicates two GSCs in a control *aub^HN2^/+* germarium (F) and the yellow dashed lines indicate differentiating cysts in the niche in *aub^HN2^/+*; *Def(Ald1)*/+ (H) and *aub^HN2^/Eno^KG01162^* (G) germaria. (I) Quantification of germaria showing GSC loss (0-1 GSC) in heterozygous and double heterozygous mutant females of the indicated genotypes, 7 days after eclosion. The number of scored germaria (n) is indicated. * *p*-value <0.05, ** *p*-value <0.01, *** *p*-value <0.001, ns, non-significant using the χ^2^ test, compared to the sum of GSC loss from the two corresponding single heterozygous. Scale bars: 10 μm.

Using a genetic approach, we analyzed the defect in GSC self-renewal when both *aub* and either *Ald1* or *Eno* gene dosage was reduced by half. The concomitant reduction of gene dosage of *aub* and its potential target *Ald1* or *Eno* led to a synergistic effect in GSC loss analyzed by immunostaining, with higher GSC loss in double heterozygous mutants than the sum of GSC loss from both heterozygous mutants (Figures 5F-5I), in lines with a positive regulation of *Ald1* and *Eno* by Aub.

Together, these results confirm the direct binding of Aub to glycolytic mRNAs and show the requirement of Aub loading for this binding. Furthermore, genetic data provide functional evidence that Aub positively regulates glycolytic mRNAs through this interaction.

### Aubergine regulation of glycolytic mRNAs depends on piRNA targeting

We then addressed the potential of piRNAs to target glycolytic mRNAs bound by Aub in GSCs, in the vicinity of GFP-Aub iCLIPs. We used recently published piRNA libraries from cultured GSCs ^49^ and analyzed potential piRNA targeting within ±60 nucleotides (nt) from significant GFP-Aub iCLIPs using various complementarities ^31^ (see Methods). We then filtered targeting with the highest piRNA occurrences (total occurrence of overlapping piRNAs >500) and a low minimum free energy prediction of the piRNA-target RNA duplex (<-24 kcal/mol) ^50^. Using these criteria, we identified four glycolytic mRNAs targeted by abundant GSC piRNAs in regions overlapping GFP-Aub iCLIPs (Figure S6A).

To functionally confirm the importance of glycolytic mRNA regulation by Aub and piRNAs for GSC maintenance, we focused on *Eno*. This gene produces different mRNAs but a single of them is highly expressed in GSCs (Figure 6A). This *Eno* mRNA is targeted in both its 5’UTR and 3’UTR by two piRNA populations produced from the *Quasimodo* transposable element (Figures 6A, S6A). We generated deletions of the piRNA target site in *Eno* 5’UTR, using the CRISPR-Cas9 approach. Two short deletions of 11 nt and 12 nt overlapping the target site of the piRNA seed region (nt 2-7) were obtained and named *Eno^Δpi11^* and *Eno^Δpi12^* (Figure 6A). Both deletions removed a donor splice site and could prevent splicing in the 5’UTR. However, we found that splicing occurred between a new downstream donor splice site and the native acceptor site (Figures 6A, S6B). We analyzed GSC loss in *Eno^Δpi11^* and *Eno^Δpi12^* mutants either homozygous or in combination with another allele, *Eno^f07543^*, using immunostaining 14 and 21 days after fly eclosion. GSC loss was significant with both deletion mutants at both time points, demonstrating that *Eno* mRNA regulation by piRNAs is required for GSC self-renewal (Figures 6B-6E).

**Figure 6.**
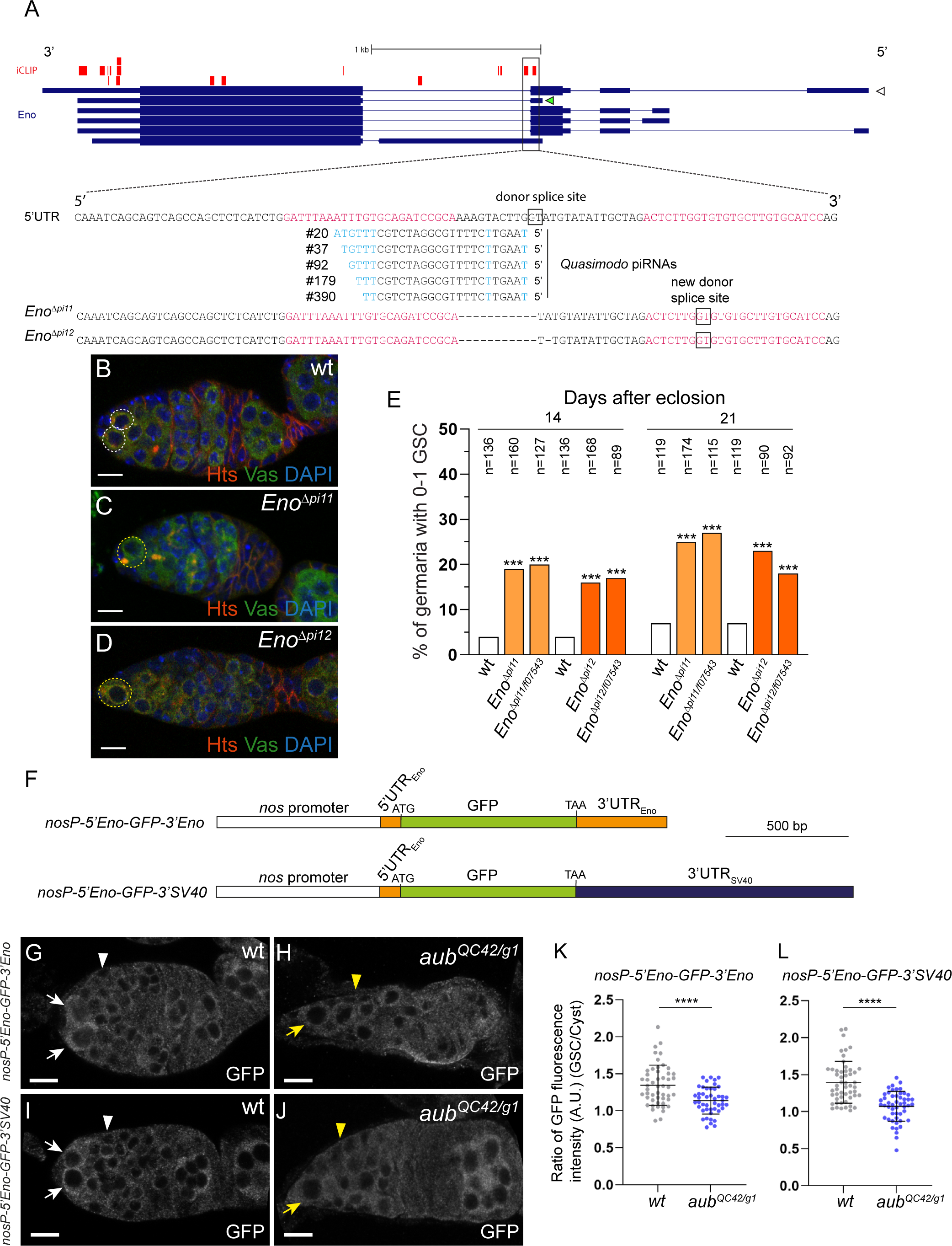
Aub-dependent activation of glycolytic mRNAs in GSCs depends on their targeting by piRNAs. (A) Schematic representation of the *Eno* gene in its genomic orientation. Thick boxes are coding sequences, thin boxes are UTRs and lines are introns. GFP-Aub iCLIPs are shown in red. iCLIPs from the three replicates performed in cultured GSCs are shown independently on three lines ^18^. The open and green arrowheads at the 5’ end of mRNAs indicate mRNA isoforms expressed in GSCs. The open arrowhead indicates low expression (>10 and <100 Transcripts Per Million (TPM): 10 to 30 TPM for this *Eno* transcript); the green arrowhead indicates high expression (>100 TPM: 150 to 240 TPM for this *Eno* transcript). The *Eno* sequence overlapping GFP-Aub iCLIPs in the 5’UTR and intron is shown. Nucleotides (nt) in red are those identified as bound by Aub in the iCLIP datasets ^18^. The sequence and occurrences of GSC *Quasimodo* piRNAs potentially targeting this region of *Eno* are indicated. Nt in blue are non-complementary nt to the *Eno* sequence. The sequences of *Eno^Δpi11^* and *Eno^Δpi12^* are shown; the deleted nt are indicated by dashes. Boxed nt indicate the donor splice site and the new donor splice site in the mutants. (B-D) Confocal images of immunostaining of wt (B), *Eno^Δpi11^* (C) and *Eno^Δpi12^* (D) germaria with anti-Vasa (green), anti-Hts (red) and DAPI (blue). The white and yellow dashed lines indicate GSCs in wt (B) and mutant (C, D) germaria, respectively. (E) Quantification of germaria showing GSC loss (0-1 GSC) in wt and mutant females, 14 and 21 days after eclosion. The number of scored germaria (n) is indicated. *** *p*-value < 0.001 using the χ^2^ test. (F) Schematic representation of *Eno* reporter transgenes. Open boxes represent *nos* promoter; orange boxes represent *Eno* 5’UTR and 3’UTR; green boxes represent the GFP coding sequence; and the blue box represents *SV40* 3’UTR. (G-J) Confocal images of immunostaining of germaria expressing *Eno* reporter transgenes *nosP-5’Eno-GFP-3’Eno* (G, H) and *nosP-5’Eno-GFP-3’SV40* (I, J), in wt (G,I) and *aub^QC42/g1^* mutant context (H, J) with anti-GFP. White and yellow arrows point to GSCs in wt and mutant germaria, respectively. White and yellow arrowheads indicate the region containing germline cysts, where GFP was quantified in wt and mutant germaria, respectively. (K,L) Quantification of GFP levels in wt and *aub^QC42/g1^* mutant GSCs and differentiating cyst cells using immunostaining experiments shown in (G-J). Fluorescence intensity was measured in arbitrary units using the ImageJ software. Ratios of fluorescence intensity of one GSC to one cyst cell per germarium were plotted. Horizontal bars represent the mean and error bars represent standard deviations. \*\*\*\**p*-value <0.0001 using the unpaired Student’s *t*-test. Scale bars: 10 μm.

We next generated *Eno* reporter transgenes containing *nanos* (*nos*) promoter to allow germline expression, followed by *Eno* 5’UTR, GFP coding sequence and either *Eno* 3’UTR or *SV40* 3’UTR (Figure 6F). Strikingly, both transgenes recapitulate Eno protein expression profile in wild-type germaria, with higher GFP levels in GSCs than in differentiating cyst cells (Figures 6G, 6I, 6K, 6L). Furthermore, this difference between GFP levels in GSCs and differentiating cells was significantly reduced in *aub* mutant germaria (Figures 6H, 6J-6L). These results show that Aub-dependent *Eno* mRNA activation depends mainly on its 5’UTR, strongly suggesting a translational activation.

These data demonstrate that mRNA activation by Aub in GSCs depends on piRNA targeting and is required for GSC self-renewal. They further show that *Eno* mRNA regulation by Aub involves its 5’UTR and likely occurs at the level of translation.

### Aub function in GSC self-renewal involves its role in activating glycolysis

Aub was shown to be essential for GSC self-renewal through its role in the regulation of cellular mRNAs ^18, 19^. We find here that Aub activates the translation of glycolytic mRNAs in GSCs and that high glycolysis is required for GSC self-renewal. Thus, we asked whether activation of glycolysis contributes to Aub function in GSC self-renewal. We analyzed if *aub* mutant phenotype of GSC loss could be rescued to some extent by increasing glycolysis. Phosphofructokinase (Pfk) catalyzes an irreversible reaction in glycolysis that corresponds to the first commitment of glucose to the glycolytic pathway. Because Pfk is rate limiting, it plays a critical role in determining glycolytic flux and is considered as the pacemaker of glycolysis ^51^. We overexpressed *Pfk* in germ cells using a *Drosophila* strain containing a *P-UASp* insertion upstream of the coding sequence, crossed with the *nos-Gal4* germline driver (Figure S7). GSC analysis using immunostaining 7 days and 14 days after fly eclosion in two different *aub* allelic combinations, revealed that GSC loss in *aub* mutant germaria was significantly reduced following germline overexpression of *Pfk* (Figures 7A-7D). In particular, the complete lack of germ cells (empty germaria) that became substantial with time due to GSC loss, sharply decreased when *Pfk* was overexpressed (Figures 7B, 7D).

**Figure 7.**
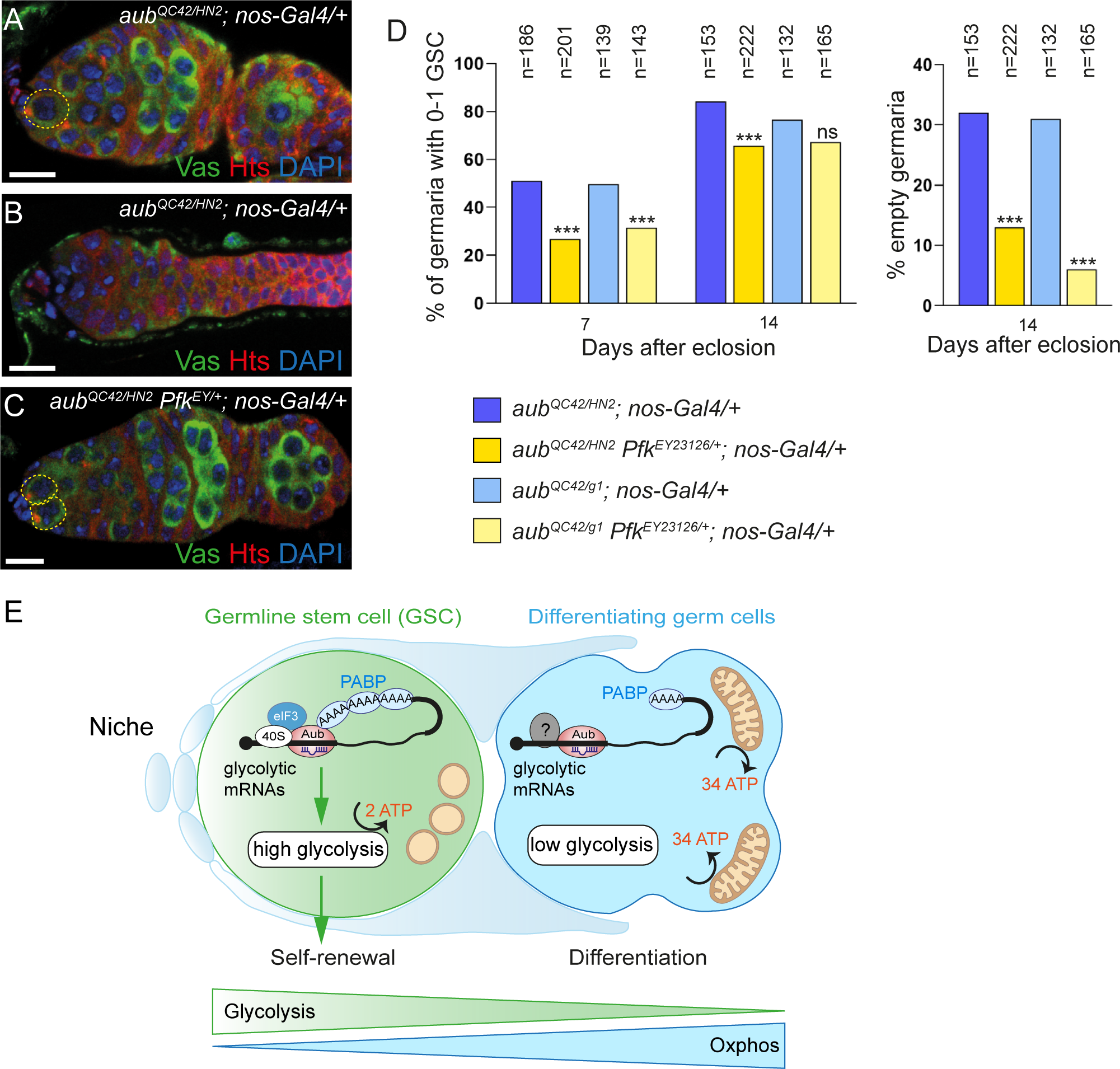
Activation of glycolysis contributes to Aub function in GSC self-renewal. (A-C) Confocal images of immunostaining of *aub^QC42/HN2^; nos-Gal4/+* (A, B) and *aub^QC42/HN2^, Pfk^EY23126/+^; nos-Gal4/+* (C) germaria with anti-Vasa (green), anti-Hts (red) and DAPI (blue). The yellow dashed lines indicate the GSCs. An empty germarium, devoid of germ cells is shown in (B). (D) Quantification of germaria showing GSC loss (0-1 GSC) in mutant females of the indicated genotypes, 7 and 14 days after eclosion, and of empty germaria in mutant females 14 days after eclosion. The number of scored germaria (n) is indicated. *** *p*-value < 0.001, ns, non-significant using the χ^2^ test. (E) Model of Aub and piRNA function in metabolic remodeling in GSCs. Aub guided by piRNAs binds glycolytic mRNAs in GSCs leading to their translational activation and higher levels of glycolytic enzymes. Translational activation might involve Aub interaction with the translation initiation factors PABP and eIF3 as is the case in the embryo ^54^. Aub-dependent activation of glycolytic mRNAs leads to high glycolysis in GSCs that is required for their self-renewal. During cyst cell differentiation, mitochondrial maturation takes place leading to an increased number of fused mitochondria equipped with cristae. This allows a gradual metabolic rewiring towards oxphos during the process of germ cell differentiation. Other factors (question mark) are expected to cooperate with Aub to restrict translational activation to GSCs. Scale bars: 10 μm.

These data provide functional evidence that Aub-dependent activation of glycolysis is a key contribution to Aub function in GSC self-renewal.

## Discussion

Over the recent years, it has become clear that, in addition to its bioenergetic functions, metabolism plays a role in the regulation of cellular programs, and in particular in the control of stem cell fate ^52^. Various populations of stem cells depend on glycolysis to generate ATP and this specific metabolism actively contributes to stemness maintenance, whereas ATP production through mitochondrial respiration is directly associated to differentiation ^6^. The molecular mechanisms of this metabolic rewiring remain poorly understood. Transcriptional regulation by HIF transcription factors are known to be involved. The stabilization of these transcription factors under hypoxic conditions in stem cell niches, leads to both transcriptional activation of glycolytic genes and reduction of mitochondrial biogenesis ^9, 10^. Another level of regulation in human pluripotent stem cells is based on increased expression of uncoupling protein 2 (UCP2) that shunts pyruvate from mitochondria, thus favoring glycolysis ^1, 53^. However, the molecular bases of this regulation remain unknown. A key information from this study is to identify an unanticipated level of control of energy metabolism in stem cells based on mRNA regulation by piRNAs and PIWI proteins. We show that Aub targets glycolytic mRNAs and this leads to elevated levels of glycolytic enzymes and higher glycolysis in GSCs. Aub loading with piRNAs is necessary for Aub interaction with glycolytic mRNAs and deleting a potential piRNA target site in *Eno* mRNA 5’UTR results in defective GSC self-renewal. Moreover, *Eno* reporter transgenes containing *Eno* UTRs with an unrelated promoter recapitulate Aub-dependent *Eno* mRNA regulation in GSCs. These results demonstrate a control of metabolic remodeling in stem cells at the posttranscriptional level involving Aub and piRNAs (Figure 7E).

The piRNA target site in *Eno* mRNA shown to be functional for *Eno* regulation in GSCs is located in the 5’UTR, 23 nt upstream of the translation initiation codon. Furthermore, an *Eno* reporter transgene containing only *Eno* 5’UTR, followed by exogenous coding sequence and 3’UTR, reproduces Aub-dependent higher protein levels in GSCs. These data point to a function of Aub and piRNAs in translational activation of glycolytic mRNAs in GSCs (Figure 7E). A role for Aub and piRNAs in translational activation has been demonstrated for *nos* mRNA in the *Drosophila* embryo^54^. It depends on direct protein interactions between Aub and the translation initiation factors poly(A) binding protein (PABP) and eIF3 subunits. Importantly, a similar role in translational activation has also been described for Miwi, the mouse homolog of Aub, and also involves direct Miwi interactions with PABP1 and subunits of eIF3 ^55^, showing that this positive function of PIWI proteins in mRNA regulation is conserved ^12^.

The function of mitochondria has been analyzed in the *Drosophila* ovarian GSC lineage, although the energetic pathways used at the different steps of GSC differentiation remain uncharacterized. Components of ATP synthase (complex V), the last complex of the mitochondrial electron transport chain, are expressed in the germarium in dividing cyst cells and required for their division and differentiation ^33, 56^. However, ATP synthase at these stages is involved in mitochondrial maturation, i.e. cristae formation, but not for oxphos. Indeed, ATP synthase requirement in differentiation of early cyst cells was shown to depend on its role in mitochondrial remodeling that prevents ER stress, premature meiosis and cell death ^56^. The formation of mitochondrial cristae increases the surface of the inner mitochondrial membrane and would accommodate the presence of more electron transport chain complexes, favoring a switch towards ATP production through oxphos ^57, 58^. However, components of complexes II, III and IV of the electron transport chain are not required for early cyst cell differentiation, indicating that early differentiating cells do not strongly depend on oxphos ^56^. The switch towards oxphos would be more gradual, and indeed mutants targeting complexes III or IV stop oogenesis after full differentiation of the egg chamber at stages 8/9. These data are in lines with the upregulation of electron transport chain complexes at the transcription level in differentiated 16 cell-cysts and with the actual rise of oxphos levels in these cells, measured through mitochondrial membrane potential ^34^.

Here, we identify the energetic pathway used in GSCs. We find that GSCs rely heavily on glycolysis to generate ATP, since reducing the levels of glycolytic enzymes leads to GSC loss. However, differentiation is not prevented in GSCs with decreased glycolysis indicating that glycolysis is not the main energetic pathway in early differentiating cyst cells. These data raise the question of which energetic pathway is used by these early cyst cells. Based on mRNA expression profiling, it was previously shown that GSCs and their immediate progeny express high levels of arginine kinase suggesting that early differentiating cyst cells might employ a phosphagen system involving arginine-phosphate and arginine kinase to use energy provided from neighboring cells ^59^. This specific metabolism could reflect the fact that selection of mitochondria for the next generation occurs in these early differentiating cyst cells ^60^. Therefore, the use of a different energetic pathway might favor this key selection process without affecting cell fate.

Consistent with the importance of glycolysis to maintain GSC stemness, we show that increasing oxphos in GSC leads to their differentiation. Similarly, interfering with mitochondrial fusion or fission in GSCs was shown to induce their differentiation by increasing oxphos ^42^.

Importantly, in *aub* mutant GSCs, the reduced levels of glycolytic enzymes and decreased lactate accumulation are accompanied by a metabolic rewiring towards oxphos. Mitochondrial maturation occurs precociously in these mutant GSCs with a large proportion of mitochondria containing cristae. Consistent with the role of ATP synthase in mitochondrial maturation, we also find increased levels of ATPsynα in *aub* mutant GSCs. Moreover, these mitochondria are active in generating ATP as measured by inner membrane potential. This metabolic switch in *aub* mutant GSCs could be indirect and reflect a metabolic adaptation to reduced glycolysis. In lines with this hypothesis, we found increased levels of ATPsynα in GSCs mutant for the Ald1 and Eno glycolytic enzymes. However, Aub does bind a large set of mRNAs ^18, 31^ and might also play a more general role in the metabolic control in the GSC lineage through the regulation of other metabolic mRNAs.

Our functional data reveal that Aub function in activating glycolysis substantially contributes to its role in GSC self-renewal, since increasing the levels of Phosphofructokinase, the rate limiting enzyme in glycolysis, significantly reduces the *aub* mutant phenotype of GSC loss. PIWI proteins are expressed in different populations of adult stem cells and were shown to be required for stem cell homeostasis. However, in most cases, the molecular mechanisms underlying this PIWI function remain uncharacterized. Since both a role of PIWI proteins in stemness and a metabolic reprogramming towards glycolysis are conserved features of stem cells, an intriguing possibility would be that the metabolic regulation by piRNAs and PIWI proteins might also be conserved in stem cells across species. In planarians, the PIWI protein SMEDWI-3 interacts with a large set of mRNAs, leading to either their decay or binding without decay ^61^. This indicates a role of SMEDWI-3 in mRNA regulation that might extend to metabolic mRNAs.

Aub has been previously described to regulate target mRNAs directly involved in stem cell homeostasis. Aub represses *Cbl* mRNA in GSCs leading to reduced protein levels, a regulation required for GSC self-renewal ^19^. In addition, Aub iCLIP in cultured GSCs identified several mRNAs encoding self-renewal and differentiation factors, among which *dunce* self-renewal mRNA was shown to be positively regulated by Aub in GSCs, whereas *bag of marbles* differentiation mRNA was activated by Aub in early differentiating cyst cells ^18^. Therefore, Aub would have a dual function in regulating mRNAs involved in both metabolism and stemness thus coupling metabolism and development to secure stem cell fate. Similarly, coregulation of both metabolism and pluripotency genes has also been reported at the transcription level through direct binding of HIF-1 to their promoters ^48^.

The importance of metabolism in piRNA biogenesis is intriguing. Both oxphos components and glycolytic enzymes play a role in piRNA production ^29, 30^. Here we show that metabolic rewiring in stem cells is regulated by Aub and piRNAs, thus revealing a crosstalk between energy metabolism and the piRNA pathway. Understanding how this crosstalk impacts on piRNA biology constitutes a major challenge for future studies.

## Acknowledgements

We are grateful to A. Arkov, T. Kai and the Bloomington *Drosophila* Stock Center for providing *Drosophila* stocks. We thank P.Y. Plaçais, T. Préat and J.R. Huynh for their gifts of plasmids and R. Lopez-Alemany for her gift of antibodies. We thank M. Jault for participating in initial experiments shown in Figure 2 and P.P. Marie for participating in experiments shown in Figures 6, S1 and S4. This work was supported by the CNRS-University of Montpellier UMR9002, ANR (ANR-15-CE12-0019-01, ANR-19-CE12-0031, ANR-21-CE12-0035-01) and FRM. PPM held a salary from FRM. CE and JC held a salary from ANR. CG held a salary from ANR and MSDAvenir, and AR held a salary from ANR and Fondation ARC.

## Methods

### Drosophila stocks

All *Drosophila* strains used in this study are listed in Table S1. *Drosophila* were raised at 25°C on standard medium. The same numbers of flies were used for control and experimental crosses and kept to a maximum of 12 females and 6 males per vial. Crosses were transferred to fresh vials every three days. All crosses involving *nos-Gal4* were performed using *nos-Gal4* females.

### DNA constructs and transgenic fly lines

To produce *UASz-Laconic* sensor, the *pUASt-Laconic* plasmid (a gift from T. Préat ^43^) was digested with *EcoR*I and *Xba*I. The resulting fragment containing the Laconic sensor coding sequence was purified from agarose gel and cloned into the *pBSII-SK* vector digested with *EcoR*I and *Xba*I. This construct was then digested with *Xho*I and the resulting fragment was purified and cloned into the *pUASz1.0* vector ^62^ digested with *Xho*I. The resulting plasmid was validated by sequencing. Transgenic lines containing *UASz-Laconic* were generated using PhiC31-based integration into the *attP-9A[VK00027]* landing site on chromosome III, by BestGene Inc. (California). *nosP-5’Eno-GFP-3’Eno* and *nosP-5’Eno-GFP-3’SV40* plasmids were generated by assembly of PCR fragments into the *pNOSPE_MCP_eGFP* vector ^63^ digested by *Not*I and *BamH*I using the NEBuilder HiFi DNA Assembly Cloning Kit (NEB). *pNOSPE_MCP_eGFP* contains the *nanos* promoter, the mini-*white* gene and an attB site for PhiC31-mediated transformation. The primers used to generate these constructs are listed in Table S2. The PCR fragments were amplified with Q5 High-Fidelity DNA Polymerase or Phusion® High-Fidelity DNA Polymerase and the resulting plasmids were validated by sequencing. Transgenic lines were generated using PhiC31-based integration into the *attP2* landing site on chromosome III, by BestGene Inc. (California).

### CRISPR/Cas9 editing

The donor plasmid *pBS-Eno-Mutpi* was generated by assembly of PCR fragments into *pBS-SK^+^* digested by *EcoR*I and *Kpn*I using the NEBuilder HiFi DNA Assembly Cloning Kit (NEB) and primers designed for amplifying ≈1 kb long homology arms from *w^1118^* genomic sequence. This donor plasmid was designed to introduce 7 point mutations in the *Quasimodo* piRNA target site in *Eno* 5’UTR. The guide RNA producing plasmid *pDCC6-Eno-gRNA* was generated as follows. *pDCC6* vector ^64^ was digested by *Bbs*I and ligated with two annealed primers corresponding to sgRNA in *Eno* 5’UTR. The primers used to generate these constructs are listed in Table S2. PCR fragments were amplified with Q5 High-Fidelity DNA Polymerase or Phusion® High-Fidelity DNA Polymerase and the resulting plasmids were validated by sequencing. The donor plasmid and *pCCD6-Eno-gRNA* that also produces the Cas9 enzyme were co-injected into the *w^1118^* stock at the Madrid *Drosophila* Transgenesis Facility (Centro de Biología Molecular Severo Ochoa). Injected flies were individually crossed to produce independent lines and these lines were screened by PCR using a primer corresponding to the mutant sequence in *Eno* 5’UTR. A CRISPR edited line containing this mutant sequence was not recovered, but two lines containing short deletions in this region were identified (*Eno*^*Δpi11*^ and *Eno*^*Δpi12*^). *Eno* sequence overlapping the donor plasmid was validated in these lines.

### Production of mitotic clones in ovaries

The Flipase under a heat shock promoter was used to induce mitotic clones in the germline of adult females. *Ald1^EY13155^*, *Eno^KG01162^, Eno^f07543^* and *aub^HN2^* mutant germline clones were induced in 3 day-old females by two 37°C heat shocks of 1 h spaced by an 8 h-recovery period at 25°C, during three consecutive days. The flies were then maintained at 25°C until ovarian dissection, either 3, 7, 14 or 21 days following the last heat shock.

### Immunostaining

Ovaries were dissected in PBS followed by a fixation in 4% paraformaldehyde in PBT (PBS supplemented with 0.1% Tween 20) for 20 min at room temperature in rotation. Ovaries were then rinsed one time for 10 min in PBT, blocked with 10% bovine serum albumin (BSA) in PBT for 1 h and incubated with primary antibodies in PBT supplemented with 1% BSA overnight at 4°C in rotation. The following primary antibodies were used at the indicated dilutions: mouse anti-Hts (1B1 clone) 1/50, rabbit anti-GFP (Invitrogen) 1/500, mouse anti-GFP (Invitrogen) 1/50, rabbit anti-Vasa (Santa Cruz Biotechnology) 1/200, rat anti-Vasa (DSHB) 1/50, rabbit anti-cleaved Caspase 3 (Cell Signaling) 1/300, rabbit anti-PyK (Sigma) 1/200, rabbit anti-Aldolase (Cell Signalling) 1/100, mouse anti-Eno (Santa Cruz Biotechnology) 1/100, and mouse anti-Atp5A (AbCam) 1/100. After incubation with primary antibodies, the ovaries were washed three times for 30 min in PBT-1% BSA and then incubated with fluorescent secondary antibodies diluted in PBT-0.1% BSA for 4 h at room temperature in rotation. After rising two times for 10 min in PBT, the ovaries were incubated in 0.1 µg/mL DAPI in PBT to stain DNA. They were then mounted in Vectashield mounting medium. Images were acquired using a Confocal Leica SP8 and analyzed with the ImageJ software. For immunostaining with anti-Atp5A, the fixation was with 5% formaldehyde in PBS for 25 min at room temperature in rotation. Ovaries were then washed with PBS followed by a permeabilization step with 1% Triton X-100 in PBS for 2 h, and blocking in PBS-1% BSA for 1 h.

### Quantification of fluorescence intensity

Confocal images of germaria were analyzed using ImageJ. The cytoplasm of one GSC and one cyst cell in a same plane were delineated using the freehand selections tool. The GFP fluorescence intensity was measured for each cell and this quantification was replicated three times. The mean of these three measurements was then calculated. The ratio: intensity in GSC/intensity in cyst cell was calculated by dividing the mean intensity value obtained for the GSC by the one obtained for the cyst cell in the same plane. The graphs were produced using the GraphPad Prism software.

### Transmission electron microscopy

The ovaries were dissected in PBS pH 7.4 and immediately fixed in 2.5% Glutaraldehyde in PHEM buffer (60 mM PIPES, 25 mM HEPES, 10 mM EGTA, 4 mM MgSO_4_·7 H_2_0) at room temperature for 1 h, and then overnight at 4°C. Ovaries were post-fixed with 1% osmium tetroxide for 1h at 4 °C and then *en bloc* stained with 1% uranyl acetate in double-distilled H_2_O at 4°C for 1 h. Dehydration series were carried with out with successive ethanol baths at 30%, 50% and 70% at 4°C, 85% at room temperature. To preserve mitochondrial crista structure, dehydration steps were limited to 5 min each. Ovaries were processed in a standard manner and embedded in Epoxy resin. 700 nm semi-thin sections were stained with 0.1% toluidine blue to evaluate the area of interest. 70 nm ultrathin sections were cut, mounted on formvar coated slotted copper grids and stained with uranyl acetate and lead citrate by standard methods. Stained grids were examined under a Tecnai G^2^ 20 S-TWIN Transmission Electron Microscope.

### Quantification of Laconic sensor FRET efficiency

Ovaries were rinsed twice in 75% ethanol and once in Schneider’s Insect Medium (Sigma) in a glass block for disinfection. They were then dissected in clean Schneider’s Insect Medium using forceps. The epithelial sheath was removed from several ovarioles using needles during no more than 20 min and ovarioles were then transferred to a drop of 20 µL of 10S oil on a high-resolution microscope slide. Confocal images and FRET efficiencies were acquired *in vivo* using a Confocal Leica SP8 microscope. Quantification of FRET efficiency was based on mTFP fluorescence before and after Venus photobleaching. The microscope settings were the following: for mTFP, excitation was at 458 nm and the detection window between 460 and 510 nm; for Venus, excitation was at 514 nm and the detection window between 560 and 620 nm. The bleaching was performed at 514 nm for 2 min. FRET efficiency was calculated as follows: (mTFP fluorescence intensity after Venus photobleaching) - (mTFP fluorescence intensity before Venus photobleaching)/(mTFP fluorescence intensity after Venus photobleaching). Fluorescence intensity was measured using ImageJ. Similar quantifications were obtained using the LEICA FRET-AB application.

### Quantification of mitochondrial membrane potential

Ovaries were dissected in Schneider’s Insect Medium (Sigma). They were stained with Deep Red FM MitoTracker (ThermoFisher Scientific) at 500 nM during 15 min at room temperature with rotation. TMRM (Biotium) was then added at 100 nM to the previous mix for 15 min at room temperature. After 3 washes in 1X PBS, ovarioles were mounted in 1X PBS on glass slides and immediately imaged using a Leica SP8 confocal microscope. Acquired images were analyzed using ImageJ. Ratiometric images were generated as follows. After inverting the colors for each channel and changing them to grey, the images were converted to 32-bit. Ratiometric images were then obtained by dividing the TMRM channel by the MitoTracker channel, using the Image Calculator function of ImageJ. The color of the resulting image was changed for Rainbow RGB. A scale bar and calibration bar were added. Quantification of mitochondrial membrane potential was performed as follows. The cytoplasm of one or two GSCs per germarium was delineated using the freehand selections tool. The fluorescence intensity of the two channels, TMRM and MitoTracker, were measured and the value obtained for the TMRM channel was divided by the value obtained for the MitoTracker channel.

### RNA-immunoprecipitations and RT-qPCR

For RNA-immunoprecipitation, ovaries from young females (1 to 3 days old) were dissected in PBS and kept at -80°C. 100 pairs of ovaries per genotype were crushed in 600 µl DXB-150 (25 mM HEPES pH 6.8, 250 mM sucrose, 1 mM MgCl_2_, 1 mM DTT, 150 mM NaCl, 0.1% Triton X-100) containing cOmplete^TM^ EDTA-free Protease Inhibitor Cocktail (Roche) and RNase Inhibitor (0.25 U/µl; Promega). 25 µl of GFP-Trap beads (ProteinTech) were incubated with the extracts for 3 h on a wheel at 4°C. The beads were then washed seven times with DXB-150 for 10 min at room temperature. RNA was prepared using TRIzol (Invitrogen), followed by DNA removal with TURBO DNA-free (Ambion). For RT-qPCR without immunoprecipitation, 10 pairs of ovaries were directly crushed in TRIzol and RNA was prepared as described above. 500 ng of total RNA was used for reverse transcription. qPCR was performed using Universal SYBR Green Master Mix and the LightCycler® 480 Instrument (Roche). Primers used for RT–qPCR are listed in Table S3.

### Bioinformatics and statistics

#### Quantification of glycolytic mRNAs using RNA-seq

We used published datasets (GSE119862) ^49^ in which we selected RNA-seq performed in cultured GSCs and *nos-Gal4/UASp-tkv^M1^* ovaries, which we analyzed using Salmon ^65^ (https://combine-lab.github.io/salmon). Transcript quantification was done using --validateMappings option with the raw data on *Drosophila melanogaster* genome (FlyBase release 6.28). Each RNA-seq dataset was analyzed independently and glycolytic mRNAs were filtered using thresholds of minimum expression of 10 and 100 Transcripts Per Million (TPM) in all replicates.

#### Construction of piRNA library from small RNA-seq in cultured GSCs

The small RNA-seq datasets in cultured GSCs (GSE119862) ^49^ were analyzed using a simplified homemade version of the pipeline piPipes ^66^ (https://github.com/bowhan/piPipes). The raw data were first trimmed with Cutadapt ^67^ (https://github.com/marcelm/cutadapt) as follow -a TGGAATTCTCGGGTGCCAAGGAACTCCAGTCACnnnnnnATCTCGTATGCCGTCTTCTGCT TG -m 18 -M 30 --discard-untrimmed. The alignment on the *Drosophila melanogaster* genome (FlyBase release 6.28) was performed using bowtie ^68^ (https://bowtie-bio.sourceforge.net/index.shtml) with the configuration -v 0 --a --best --strata. All mapped reads were next annotated in the following order to rRNAs, snoRNAs, tRNAs, snRNAs, small mtRNAs and miRNAs using bowtie with the same parameters as above, except for rRNAs where only the best alignment was reported (-m 1). At each annotation step, unmapped reads were conserved (--un option) and used as input against the next small RNA species. All non-coding RNA references were obtained from diverse sources : NCBI (https://www.ncbi.nlm.nih.gov), FlyBase (http://flybase.org) and RNAcentral (https://rnacentral.org). The remaining reads were then trimmed by size, selecting only the 23-29 nt, and the pool was cleaned by collapsing identical sequences and attributing a unique identifier to each, along with the number of times a read had been retrieved in the whole pool. We got a library of 4,370,902 sequences of putative piRNAs. The library was then annotated by independent mappings against transposable elements and piRNA clusters (from piPipes) and *Drosophila* transcriptome (from FlyBase), with the following bowtie’s options (transposons : -v 3 -a --best --strata; piRNA clusters : -v 0 -m 1; transcriptome : -v 0 -m 1 --norc). Using a handmade script, annotations were next merged with sequence headers alongside the unique identifier. Unannotated piRNAs were kept and their annotation field complemented with a « 0 ».

#### piRNA targeting close to Aub binding sites in glycolytic mRNAs

Aub cross-link sites in cultured GSCs were obtained from published iCLIP experiments (GSE96751) ^18^ and iCLIP genomic coordinates were converted to the dm6 release using the online converter of FlyBase. Using our piRNA library, bowtie was configured with different settings to identify potential target sites on transcripts. Nine targeting conditions were defined as follows: 0 to 3 mismatches within the piRNA sequence using option -v set up from 0 to 3; targeting of a 5’ seed of 16 nt without mismatch, 18 nt with 0 to 2 mismatches, or 20 nt with 1 mismatch using options -l and -n. For all targeting conditions, the first nt of the piRNA was excluded from the alignment using the -5 option. Each targeting was done on the whole *Drosophila* genome. All valid alignments were reported and when a seed was defined, the -e option was set to an arbitrary value of 2000, disabling the quality values. Using the BEDtools Suite ^69^ (https://bedtools.readthedocs.io/en/latest/content/bedtools-suite.html), the alignment files were next converted into the bed format and piRNA coordinates were overlapped with protein coding genes coordinates obtained from FlyBase. Only antisense hits were reported using the –wa -wb -S options of intersectBED. Finally, antisense piRNAs closed to Aub cross-link sites were conserved using windowBED with a bin size of 60 nt in each direction of a cross-link site (-r 60 –l 60). Targeting results were then visualized and studied on the Integrative Genomics Viewer ^70^ (https://software.broadinstitute.org/software/igv).

#### Calculation of MFE scores

We evaluated each piRNA-mRNA duplexes by calculating the Minimum Free Energy (MFE) using the RNAduplex tool from the ViennaRNA package ^71^ (https://www.tbi.univie.ac.at/RNA/RNAduplex.1.html).

#### Statistical tests

Statistical tests were performed using GraphPad Prism or online χ^2^ test (http://www.aly-abbara.com/utilitaires/statistiques/khi_carre.html) softwares.

