## Supplemental Figures for "piRNAs are regulators of metabolic reprogramming in stem cells"

### Extended data

#### Extended data Figure legends

**Figure S1. Expression and function of glycolytic enzymes in germaria.**

(A-F) Specificity of antibodies against glycolytic enzymes in immunostaining. Confocal images of immunostaining of control *nos-Gal4/+* germaria (A, C, E) and germaria expressing RNAi against glycolytic mRNAs in germ cells (*UAS-Ald1-RNAi/+; nos-Gal4/+* (B), *nos-Gal4/UAS-Eno-RNAi* (D), *nos-Gal4/UAS-PyK-RNAi* (F)), with anti-Ald (A, B), anti-Eno (C, D) and anti-PyK (E,F) showing the reduced staining intensity in germ cells, in the presence of RNAi.

(G-H) Quantification of GFP-PyK fusion protein in GSCs and differentiating cyst cells. GFP-PyK is produced from the *Wee-P338* stock that contains a *P-GFP* insertion within the *PyK* gene [1]. Confocal images of immunostaining of *Wee-P338* germaria with anti-GFP (green), anti-Hts (red) and DAPI (blue) (G, G'). White arrows indicate the GCSs and white arrowheads indicate the region containing germline cysts, where glycolytic enzyme levels were quantified. Quantification of GFP protein levels in GSCs and differentiating cyst cells (H). Fluorescence intensity was measured in arbitrary units using the ImageJ software. Horizontal bars represent the mean and error bars represent standard deviations. $^{**}p$-value $<0.01$ using the paired Student's *t*-test.

(I) Schematic representation of *Eno* and *Ald1* mutants used in this study. Open boxes represent coding sequences, black boxes represent UTRs and lines represent introns. The mutations correspond to engineered *P*-element insertions and are indicated in red.

(J-N) Clonal *Ald1* and *Eno* mutant GSCs are not lost by apoptosis. Confocal images of mosaic germaria containing control (J, J', L, L'), *Ald1*$^{EY13155}$ (K, K') or *Eno*$^{KG01162}$ (M, M') clonal cells stained with anti-GFP (green) (lack of GFP indicates clonal cells), anti-cleaved Caspase 3 (red) and DAPI (blue), 3 days after clonal induction. White and yellow arrows indicate control and mutant GSCs, respectively. Red arrowheads indicate somatic cells showing cleaved Caspase 3 signal as staining positive control. Quantification of germaria with cleaved Caspase 3 positive GSCs, 3 days and 7 days after clonal induction (ACI), showing that mutant GSCs do not contain cleaved Caspase 3 more frequently than control GSCs (N). The number of scored germaria is indicated (n). **p*-value <0.05, ns, non-significant using the $\chi^2$ test.

Scale bars: 10 μm.

**Figure S2. Glycolytic genes are required for GSC self-renewal.**

(A) Quantification of glycolytic mRNAs using RT-qPCR in control ovaries (*nos-Gal4/+*) and ovaries expressing RNAi in germ cells with the *nos-Gal4* driver from 7 day old-females. mRNA levels were normalized to *sop* mRNA and set to 1 in control *nos-Gal/+* ovaries. Mean of two to three biological replicates quantified in triplicates. Error bars represent standard deviation. **p*-value <0.05, ***p*-value <0.01, ****p*-value <0.001, *****p*-value <0.0001 using the unpaired Student's *t*-test.

(B-K) GSC loss following expression of RNAi against glycolytic mRNAs in germ cells. Confocal images of immunostaining of control germaria (*nos-Gal4/+*) and germaria expressing RNAi in germ cells from 7 day old-females with anti-Hts (red) and DAPI (blue). White dashed line in (B) indicate two GSCs in a control (*nos-Gal4/+*) germarium and yellow dashed lines in (C-K) indicate a single GSC per germarium or a differentiated cyst in the niche.

(L) Quantification of germaria with GSC loss (0-1 GSC) 7, 14 and 21 days after eclosion. The number of scored germaria (n) is indicated**.** *$p$-value <0.05, **$p$-value <0.01, ***$p$-value <0.001, ns, non-significant using the $\chi^2$ test.

(M-P) Quantification of *Pdp, Pdk, ewg and spargel* mRNAs using RT-qPCR in control ovaries (*nos-Gal4/+*) and ovaries expressing RNAi or mRNA downstream *P-UAS* transgenes in germ cells with the *nos-Gal4* driver from 3 day old-females. mRNA levels were normalized to *sop* mRNA and set to 1 in control *nos-Gal/+* ovaries. Mean of two to three biological replicates quantified in triplicates or duplicates. Error bars represent standard deviation. *$p$-value <0.05, **$p$-value <0.01, ****$p$-value <0.0001 using the unpaired Student's $t$-test.

Scale bars: 10 μm.

**Figure S3. Venus alone does not produce FRET signal.**

(A-B') Confocal images of *nos-Gal4/UASp-Krimper-Venus* germaria before and after photobleaching in the white square, showing the absence of false FRET signal by photoconversion of Venus into CFP-like species emitting at 460-510 nm following photobleaching. Krimper-Venus was used as a Venus fusion protein expressed in GSCs and germ cells.

Scale bar: 10 μm.

**Figure S4. Clonal *Ald1* and *Eno* mutant GSCs express higher levels of ATPsynα.**

(A-B', D-E') Confocal images of mosaic germaria containing wild-type (A, A', D, D'), $Ald1^{EY13155}$ (B, B') or $Eno^{f07543}$ (E, E') clonal GSCs, stained with anti-GFP (green), anti-ATPsynα (red) and DAPI (blue). White arrows indicate non-clonal ($GFP^+$) and clonal ($GFP^-$) GSCs in wild-type germaria and non-clonal GSCs in mutant germaria. Yellow arrows indicate clonal $Adl1^{EY13155}$ or $Eno^{f07543}$ mutant GSCs.

(C, F) Quantification of ATPsynα protein levels in wild-type non-clonal ($GFP^+$) and clonal ($GFP^-$) GSCs in control and mutant germaria using fluorescence intensity of immunostaining with anti-ATPsynα. Fluorescence intensity was measured in arbitrary units using the ImageJ software. Horizontal bars represent the mean and error bars represent standard deviations. **p*-value <0.05, ****p*-value <0.001, ns: non-significant using the paired Student's *t*-test.

Scale bars: 10 μm.

**Figure S5. Aub binds glycolytic mRNAs in GSCs.**

(A) Schematic representation of glycolytic genes found in GFP-Aub iCLIP datasets from GSCs [2]. Thick boxes are coding sequences, thin boxes are UTRs and lines are introns. GFP-Aub iCLIPs are shown in red. iCLIPs from the three replicates performed in cultured GSCs are shown independently on three lines [2]. Open and green arrowheads at the mRNA 5' end indicate which mRNA isoforms are expressed in GSCs using RNA-seq datasets from two conditions, cultured GSCs and *nos-Gal4/UASp-tkv*$^{M1}$ ovaries [3]. Open arrowheads indicate low expression (>10 and <100 TPM in each RNA-seq replicate) and green arrowheads indicate high expression (>100 TPM in each RNA-seq replicate).

(B) Quantification of mRNAs using RT-qPCR in GFP-Aub and GFP-Aub$^{AA}$ IPs in ovaries shown in Figure 5C. mRNA enrichment was calculated compared to mRNA levels in $w^{1118}$ IPs. mRNA levels were normalized to U1 snRNA. *nos* mRNA was used as a positive control [4]. Mean of two biological replicates. mRNA enrichment was set to 1 in GFP-Aub$^{AA}$ IP. Error bars represent standard deviation.

**Figure S6. Targeting of glycolytic mRNAs by GSC piRNAs.**

(A) Schematic representation of glycolytic genes bound by GFP-Aub in GSCs. Thick boxes are coding sequences, thin boxes are UTRs and lines are introns. GFP-Aub iCLIPs are shown in red and the three replicates performed in cultured GSCs are shown independently on three lines [2]. Open and green arrowheads at the mRNA 5' end indicate which mRNA isoforms are expressed in GSCs using RNA-seq datasets from two conditions, cultured GSCs and *nos-Gal4/UASp-tkv*$^{M1}$ ovaries [3]. Open arrowheads indicate low expression (>10 and <100 TPM in each RNA-seq replicate) and green arrowheads indicate high expression (>100 TPM in each RNA-seq replicate). The sequences of the boxed regions and of the potentially targeting piRNAs are shown. Nt in red are those identified as bound by Aub in the iCLIP datasets [2]. Nt in blue are non-complementary nt, and nt in green correspond to the initiation codon in *Gapdh1* mRNA. Occurrences of piRNAs in GSCs are indicated, as well as the total of all GSC piRNAs targeting the same region when some piRNAs are not represented [3]. The minimum free energy predictions of piRNA-target RNA duplexes are indicated in yellow [kcal/mol]. Note that GFP-Aub iCLIP reads in *Gapdh1* mRNA did not reach the threshold to consider *Gapdh1* as significantly bound by Aub [2]. However, *Gapdh1* is represented because it is potentially targeted by a high number of GSC piRNAs and we identified a ping-pong signature: *Gapdh1* piRNAs overlapping over 10 nt piRNAs in opposite orientation, suggesting the actual targeting of *Gapdh1* by these piRNAs.

(B) Splicing of *Eno* short mRNA isoform in $Eno^{\Delta pi11}$ and $Eno^{\Delta pi12}$ mutants. The 5' region of this *Eno* mRNA is represented. The thick box is the coding sequence, thin boxes are UTRs and the line is the intron. The wild-type sequence and those of $Eno^{\Delta pi11}$ and $Eno^{\Delta pi12}$ mutants are shown. Nt in red are those identified as bound by Aub in the iCLIP datasets. The intronic sequence is in lower case letters. The initiation codon is in green. The deleted nt in the mutants are indicated by dashes. Donor and acceptor splice sites are boxed. The new splicing in $Eno^{\Delta pi11}$ and $Eno^{\Delta pi12}$ mutants was determined by sequencing.

**Figure S7. Overexpression of *Pfk* in germ cells.**

Quantification of *Pfk* mRNA using RT-qPCR in control ovaries (*nos-Gal4/+*) and ovaries expressing *Pfk* mRNAs downstream of a *P-UAS* transgene in germ cells ($Pfk^{EY23126}$) with the *nos-Gal4* driver, from 3 day old-females. mRNA levels were normalized to *sop* mRNA and set to 1 in control *nos-Gal/+* ovaries. Mean of three biological replicates quantified in triplicates. Error bars represent standard deviation. *$p$-value <0.05, using the unpaired Student's $t$-test.

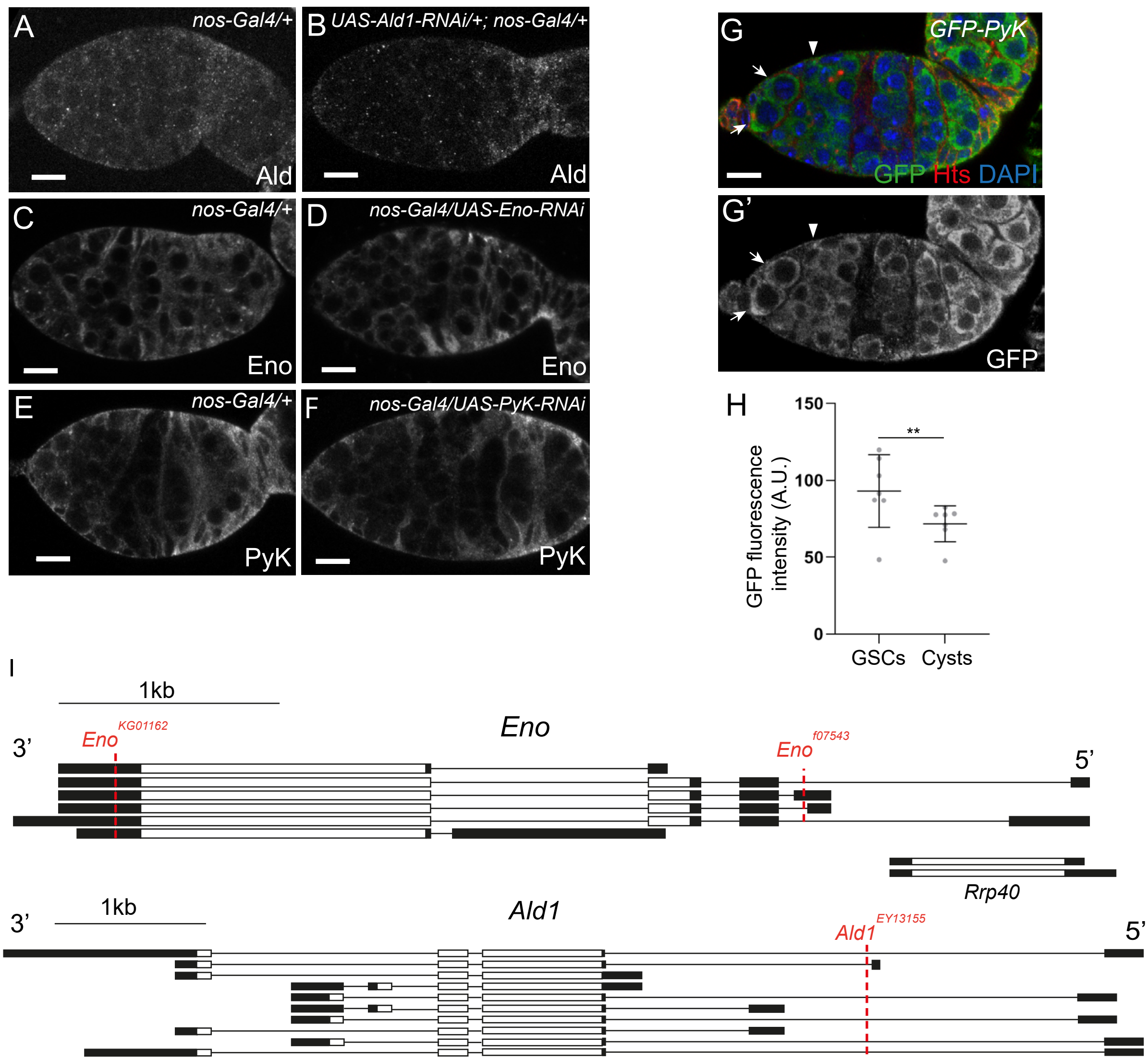


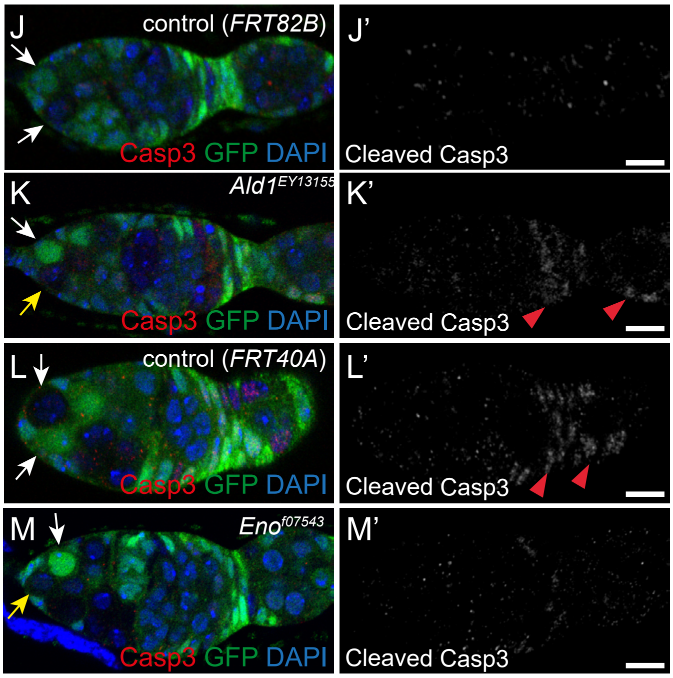


N

% of germaria with cleaved-Casp3 positive GSCs

| Genotype | 3 days ACI | | 7 days ACI | |
|---|---|---|---|---|
| Control (*FRT82B*) | 0.7 % (n=138) | ns | 5 % (n=78) | * |
| *Ald1*$^{EY13155}$ | 4.5 % (n=45) | | 1 % (n=82) | |
| Control (*FRT40A*) | 0 % (n=33) | ns | 1 % (n=84) | ns |
| *Eno*$^{f07543}$ | 3 % (n=68) | | 3 % (n=69) | |

Figure S1

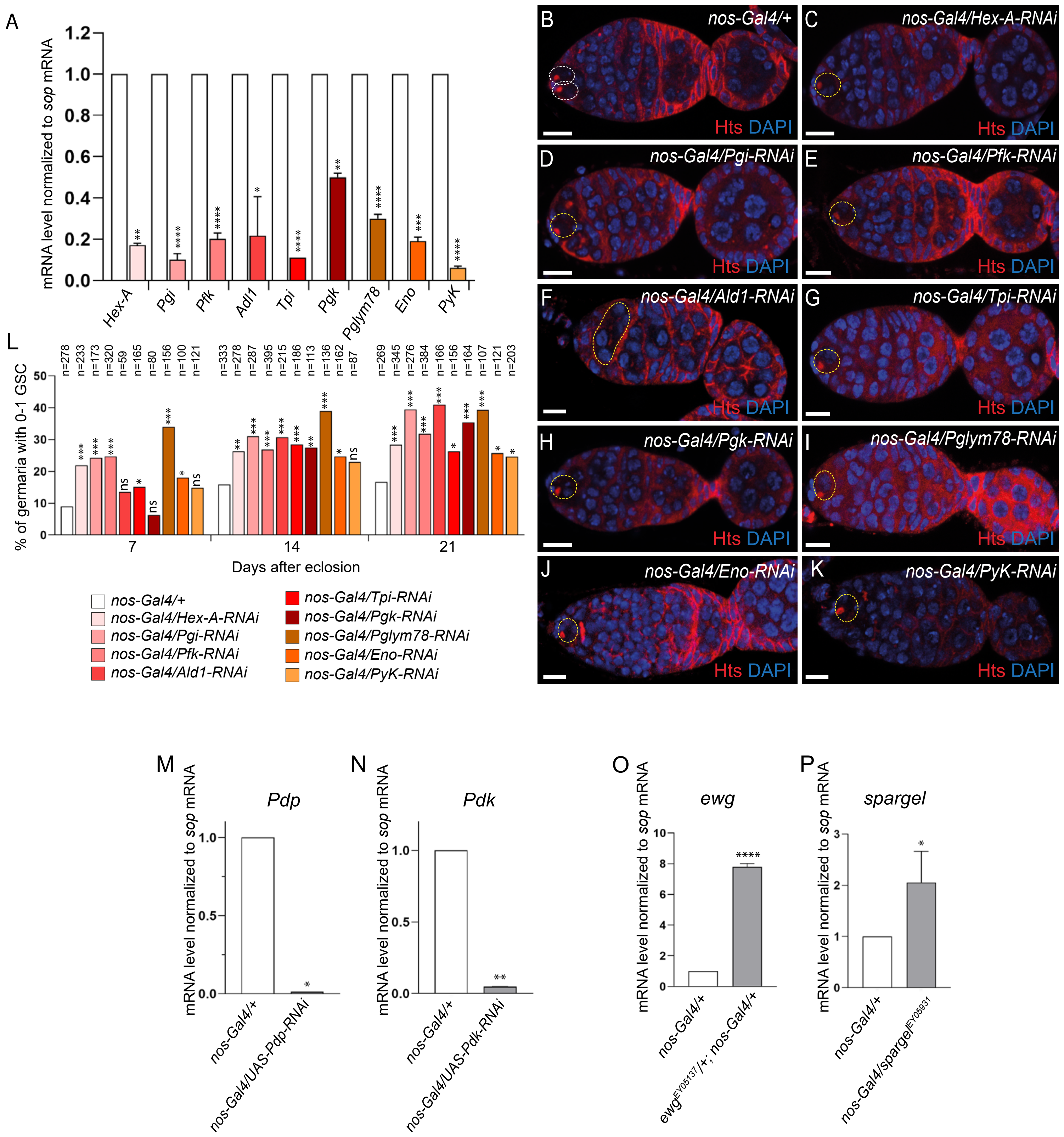
A
mRNA level normalized to sop mRNA
Hex-A
Pgi
Pfk
Adl1
Tpi
Pgk
Pglym78
Eno
PyK
L
% of germaria with 0-1 GSC
Days after eclosion
nos-Gal4/+
nos-Gal4/Hex-A-RNAi
nos-Gal4/Pgi-RNAi
nos-Gal4/Pfk-RNAi
nos-Gal4/Ald1-RNAi
nos-Gal4/Tpi-RNAi
nos-Gal4/Pgk-RNAi
nos-Gal4/Pglym78-RNAi
nos-Gal4/Eno-RNAi
nos-Gal4/PyK-RNAi
B nos-Gal4/+
C nos-Gal4/Hex-A-RNAi
D nos-Gal4/Pgi-RNAi
E nos-Gal4/Pfk-RNAi
F nos-Gal4/Ald1-RNAi
G nos-Gal4/Tpi-RNAi
H nos-Gal4/Pgk-RNAi
I nos-Gal4/Pglym78-RNAi
J nos-Gal4/Eno-RNAi
K nos-Gal4/PyK-RNAi
Hts DAPI
M Pdp
nos-Gal4/UAS-Pdp-RNAi
N Pdk
nos-Gal4/UAS-Pdk-RNAi
O ewg
ewgEY05137/+; nos-Gal4/+
P spargel
nos-Gal4/spargelEY05931

Figure S2

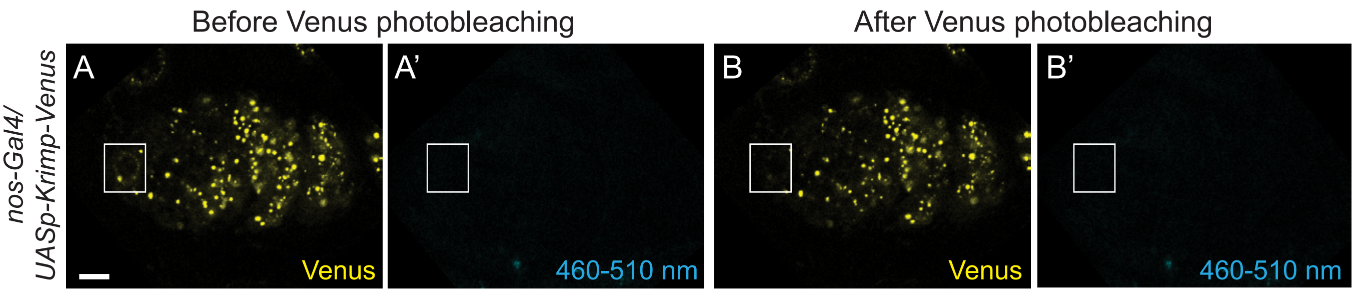
Before Venus photobleaching
After Venus photobleaching
nos-Gal4/
UASp-Krimp-Venus
A
A'
B
B'
Venus
460-510 nm
Venus
460-510 nm

Figure S3

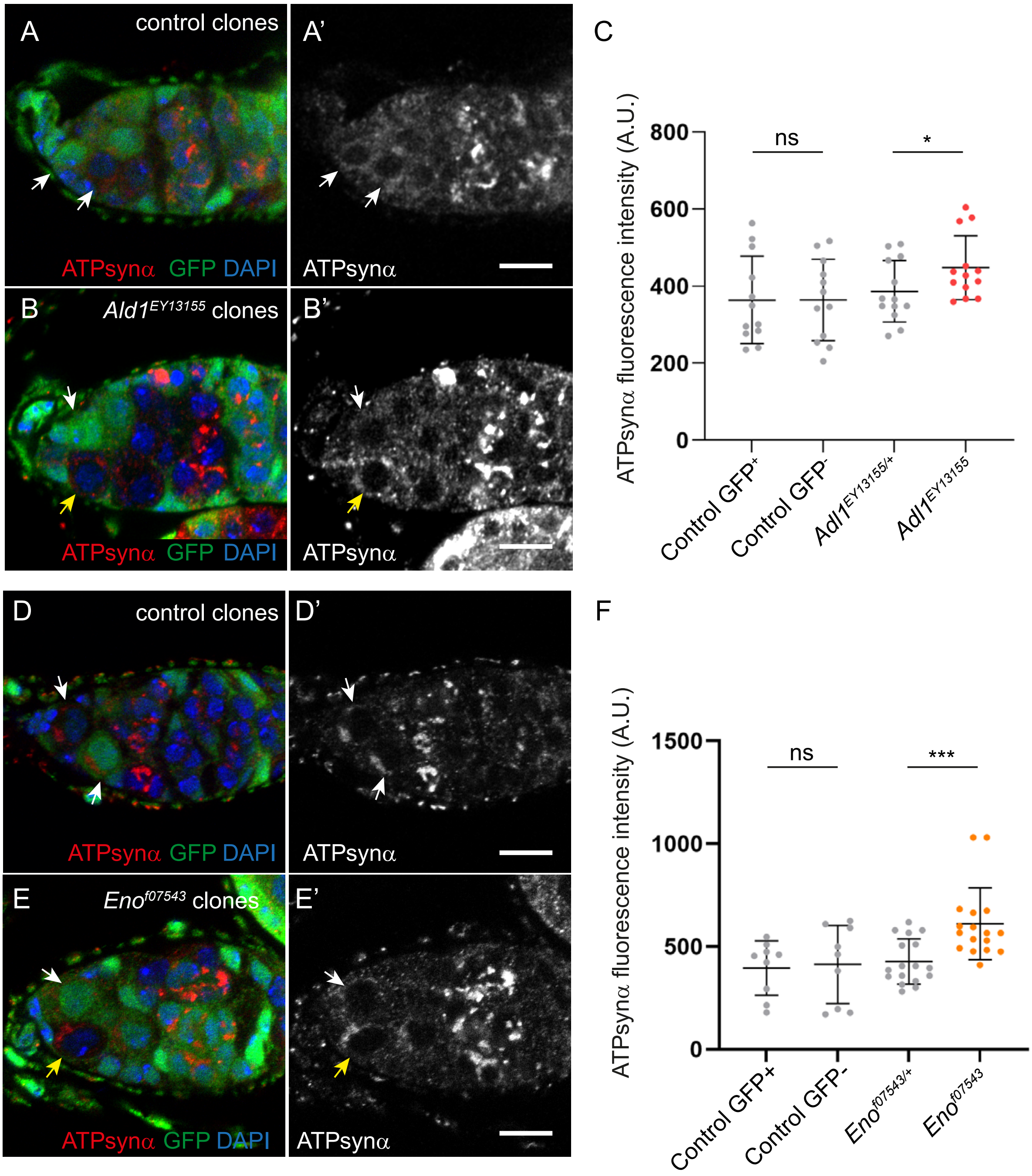
A
control clones
A'
ATPsynα GFP DAPI
ATPsynα
B
Ald1EY13155 clones
B'
ATPsynα GFP DAPI
ATPsynα
C
800
600
400
200
0
ns
*
ATPsynα fluorescence intensity (A.U.)
Control GFP+
Control GFP-
Adl1EY13155/+
Adl1EY13155
D
control clones
D'
ATPsynα GFP DAPI
ATPsynα
E
Enof07543 clones
E'
ATPsynα GFP DAPI
ATPsynα
F
1500
1000
500
0
ns
***
ATPsynα fluorescence intensity (A.U.)
Control GFP+
Control GFP-
Enof07543/+
Enof07543

Figure S4

A

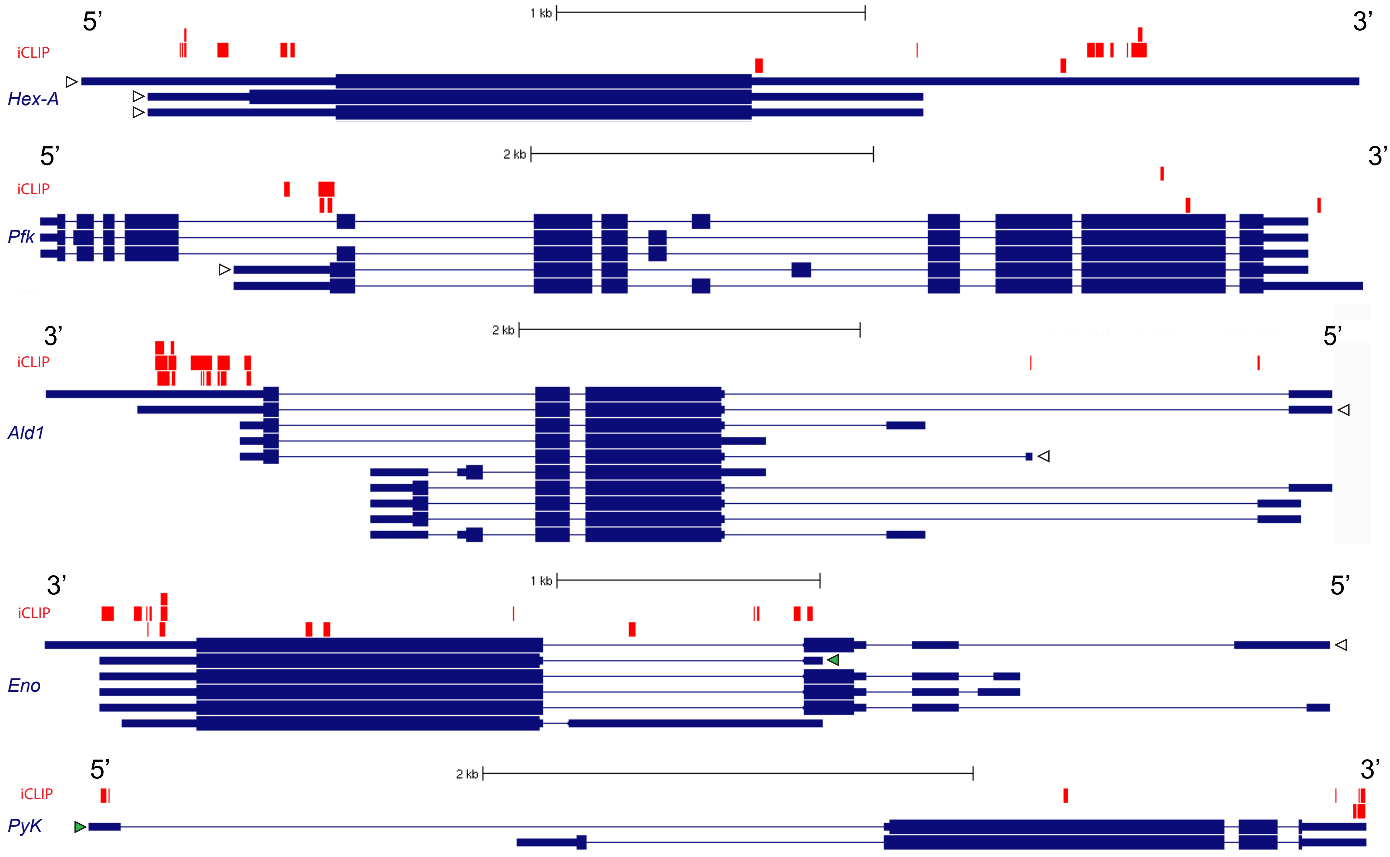

5'
3'
1 kb
iCLIP
Hex-A
2 kb
Pfk
Ald1
Eno
PyK


B

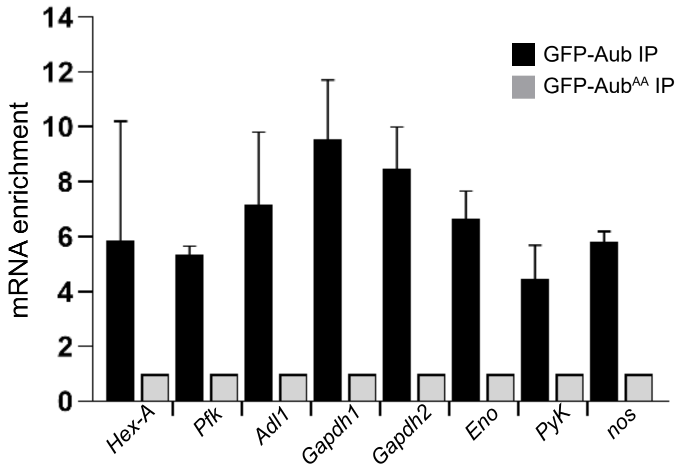

mRNA enrichment
14
12
10
8
6
4
2
0
GFP-Aub IP
GFP-Aub$^{AA}$ IP
Hex-A
Pfk
Adl1
Gapdh1
Gapdh2
Eno
PyK
nos


Figure S5

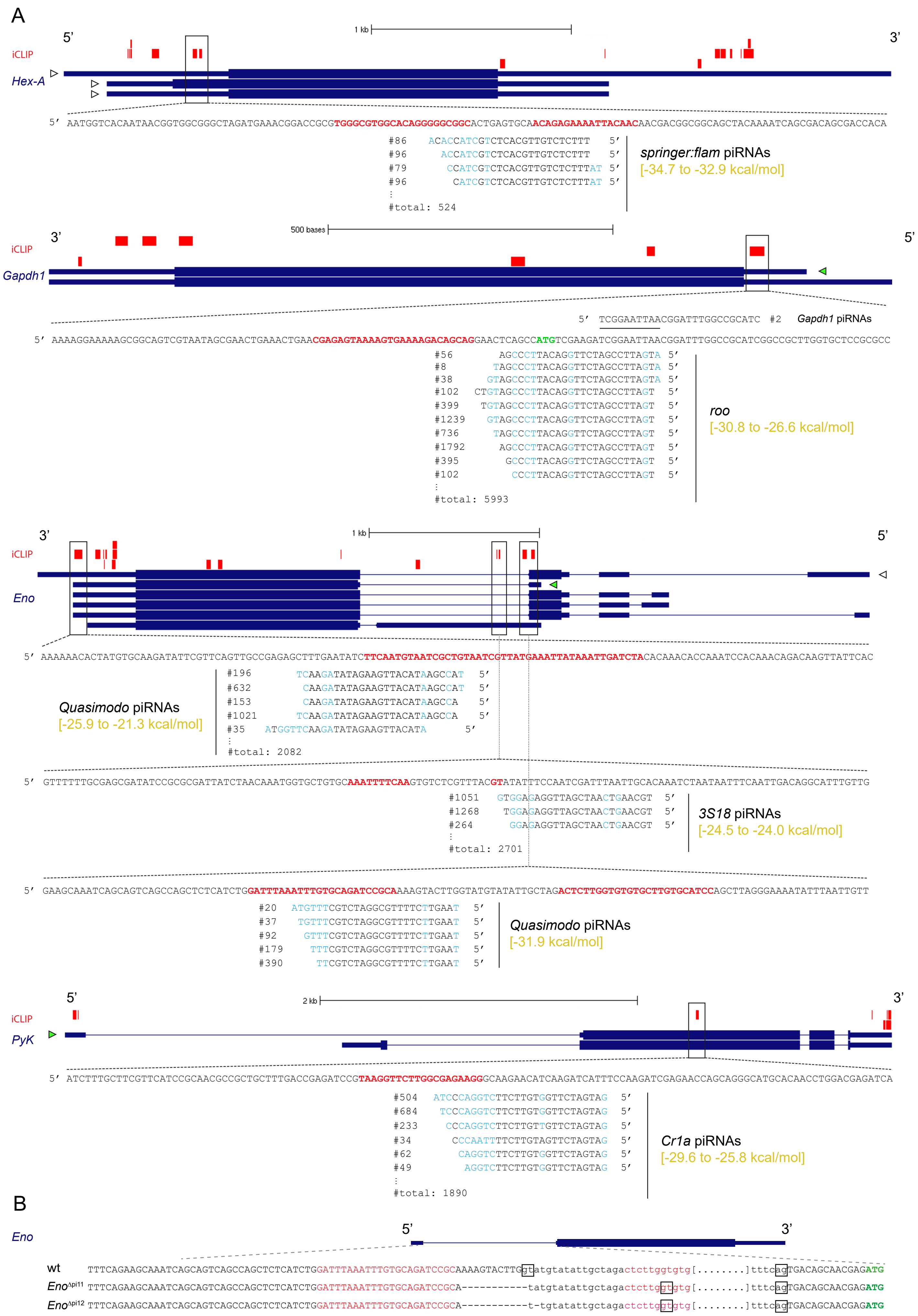

A
iCLIP
Hex-A
1 kb
5'
3'
5' AATGGTCACAATAACGGTGGCGGGCTAGATGAAACGGACCGCGTGGGCGTGGCACAGGGGGCGGCACTGAGTGCAACAGAGAAAATTACAACAACGACGGCGGCAGCTACAAAATCAGCGACAGCGACCACA
#86 ACACCATCGTCTCACGTTGTCTCTTT 5'
#96 ACCATCGTCTCACGTTGTCTCTTT 5'
#79 CCATCGTCTCACGTTGTCTCTTTAT 5'
#96 CATCGTCTCACGTTGTCTCTTTAT 5'
#total: 524
springer:flam piRNAs
[-34.7 to -32.9 kcal/mol]
iCLIP
Gapdh1
500 bases
3'
5'
5' TCGGAATTAACGGATTTGGCCGCATC #2
Gapdh1 piRNAs
5' AAAAGGAAAAAGCGGCAGTCGTAATAGCGAACTGAAACTGAACGAGAGTAAAAGTGAAAAGACAGCAGGAACTCAGCCATGTCGAAGATCGGAATTAACGGATTTGGCCGCATCGGCCGCTTGGTGCTCCGCGCC
#56 AGCCCTTACAGGTTCTAGCCTTAGTA 5'
#8 TAGCCCTTACAGGTTCTAGCCTTAGTA 5'
#38 GTAGCCCTTACAGGTTCTAGCCTTAGTA 5'
#102 CTGTAGCCCTTACAGGTTCTAGCCTTAGT 5'
#399 TGTAGCCCTTACAGGTTCTAGCCTTAGT 5'
#1239 GTAGCCCTTACAGGTTCTAGCCTTAGT 5'
#736 TAGCCCTTACAGGTTCTAGCCTTAGT 5'
#1792 AGCCCTTACAGGTTCTAGCCTTAGT 5'
#395 GCCCTTACAGGTTCTAGCCTTAGT 5'
#102 CCCTTACAGGTTCTAGCCTTAGT 5'
#total: 5993
roo
[-30.8 to -26.6 kcal/mol]
iCLIP
Eno
1 kb
3'
5'
5' AAAAAACACTATGTGCAAGATATTCGTTCAGTTGCCGAGAGCTTTGAATATCTTCAATGTAATCGCTGTAATCGTTATGAAATTATAAATTGATCTACACAAACACCAAATCCACAAACAGACAAGTTATTCAC
#196 TCAAGATATAGAAGTTACATAAGCCAT 5'
#632 CAAGATATAGAAGTTACATAAGCCAT 5'
#153 CAAGATATAGAAGTTACATAAGCCA 5'
#1021 TCAAGATATAGAAGTTACATAAGCCA 5'
#35 ATGGTTCAAGATATAGAAGTTACATA 5'
#total: 2082
Quasimodo piRNAs
[-25.9 to -21.3 kcal/mol]
5' GTTTTTTGCGAGCGATATCCGCGCGATTATCTAACAAATGGTGCTGTGCAAATTTTCAAGTGTCTCGTTTACGTATATTTCCAATCGATTTAATTGCACAAATCTAATAATTTCAATTGACAGGCATTTGTTG
#1051 GTGGAGAGGTTAGCTAACTGAACGT 5'
#1268 TGGAGAGGTTAGCTAACTGAACGT 5'
#264 GGAGAGGTTAGCTAACTGAACGT 5'
#total: 2701
3S18 piRNAs
[-24.5 to -24.0 kcal/mol]
5' GAAGCAAATCAGCAGTCAGCCAGCTCTCATCTGGATTTAAATTTGTGCAGATCCGCAAAAGTACTTGGTATGTATATTGCTAGACTCTTGGTGTGTGCTTGTGCATCCAGCTTAGGGAAAATATTTAATTGTT
#20 ATGTTTCGTCTAGGCGTTTTCTTGAAT 5'
#37 TGTTTCGTCTAGGCGTTTTCTTGAAT 5'
#92 GTTTCGTCTAGGCGTTTTCTTGAAT 5'
#179 TTTCGTCTAGGCGTTTTCTTGAAT 5'
#390 TTCGTCTAGGCGTTTTCTTGAAT 5'
Quasimodo piRNAs
[-31.9 kcal/mol]
iCLIP
PyK
2 kb
5'
3'
5' ATCTTTGCTTCGTTCATCCGCAACGCCGCTGCTTTGACCGAGATCCGTAAGGTTCTTGGCGAGAAGGGCAAGAACATCAAGATCATTTCCAAGATCGAGAACCAGCAGGGCATGCACAACCTGGACGAGATCA
#504 ATCCCAGGTCTTCTTGTGGTTCTAGTAG 5'
#684 TCCCAGGTCTTCTTGTGGTTCTAGTAG 5'
#233 CCCAGGTCTTCTTGTTGTTCTAGTAG 5'
#34 CCCAATTTTCTTGTAGTTCTAGTAG 5'
#62 CAGGTCTTCTTGTGGTTCTAGTAG 5'
#49 AGGTCTTCTTGTGGTTCTAGTAG 5'
#total: 1890
Cr1a piRNAs
[-29.6 to -25.8 kcal/mol]
B
Eno
5'
3'
wt TTTCAGAAGCAAATCAGCAGTCAGCCAGCTCTCATCTGGATTTAAATTTGTGCAGATCCGCAAAAGTACTTGgtatgtatattgctagactcttggtgtg[........]tttcagTGACAGCAACGAGATG
Eno^Api11 TTTCAGAAGCAAATCAGCAGTCAGCCAGCTCTCATCTGGATTTAAATTTGTGCAGATCCGCA-----------tatgtatattgctagactcttggtgtg[........]tttcagTGACAGCAACGAGATG
Eno^Api12 TTTCAGAAGCAAATCAGCAGTCAGCCAGCTCTCATCTGGATTTAAATTTGTGCAGATCCGCA-----------t-tgtatattgctagactcttggtgtg[........]tttcagTGACAGCAACGAGATG


Figure S6

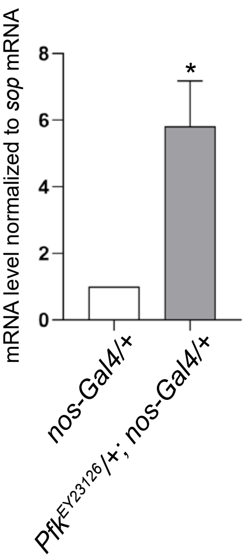
mRNA level normalized to sop mRNA
8
6
4
2
0
*
nos-Gal4/+
Pfk^EY23126/+; nos-Gal4/+

Figure S7
